# Morpho-functional timeline of progressive cystic fibrosis pancreatic exocrine and endocrine pathology derived from semi-quantitative scoring and AI-driven quantitative image analysis

**DOI:** 10.1101/2025.01.14.631729

**Authors:** Yara Al-Selwi, Dina Tiniakos, Sarah J Richardson, Christine S Flaxman, Lydia Russell, Matthew Palmer, Rowan Coulthard, Rashmi Maheshwari, Nicola Dyson, Minna Honkanen-Scott, Günter Klöppel, James AM Shaw, Nicole Kattner

## Abstract

Cystic fibrosis (CF) is associated with pancreatic exocrine insufficiency (PEI) early in life and diabetes in up to 50% of adults. The underlying CF-related sequential changes within the pancreas associated with exocrine and endocrine insufficiency remain incompletely understood due to scarcity of available human tissue, protracted disease course and absence of established robust and reproducible analytical approaches. This study aimed to develop and apply a systematic analysis cross-sectionally to CF pancreatic tissue samples from donors over a wide age range to construct a timeline related to the main exocrine and endocrine changes underlying progressive disease. Based on a histopathological semi-quantitative scoring system and AI-driven quantitative image analysis pancreatic changes were individually evaluated and classified according to three patterns: fibrotic; fibrotic and lipotic; and lipoatrophic. This systematic evaluation was applied to 29 CF and 58 control donors without pancreatic disease. Rapid loss of acinar tissue with virtually complete absence by the age of 7 years was confirmed, mirrored by fatty tissue replacement – changes underlying PEI and likely preceding progression towards diabetes. Ductal blockage by thickened secretions was associated with increasing ductal dilatation accompanied by peri-ductal fibrosis, followed by ductal loss with involution of associated fibrosis in parallel with increasing adipocyte proportional area (PA). Remaining ducts were relatively small surrounded by residual fibrosis. Islets became increasingly clustered initially surrounded by activated pancreatic stellate cells (PSCs) and fibrosis and then disorganised by interposing fibrotic tissue between endocrine cell regions and surrounded by residual collagen stranding in a ‘lipoatrophic’ pancreas. Overall islet mass was not significantly reduced but β-cell PA was significantly reduced from birth without further loss over time. We concluded that the natural history of pancreatic CF progresses inexorably from peri-ductal fibrosis to global fat replacement with relatively well-maintained islet mass but PSC-associated fibrotic islet remodelling circumstantially implicated in β-cell failure.

## Introduction

Cystic fibrosis (CF) is a genetic disease with autosomal recessive inheritance that affects multiple organs and progresses to complete functional loss of the pancreas. It is caused by a mutation in the gene encoding the CF transmembrane conductance regulator (CFTR) leading to absence or impaired function of this epithelial membrane bicarbonate and chloride transporter (Elborn, 2016). This results in accumulation of viscous mucin in the airways, gastrointestinal and reproductive tracts. In the pancreas, CF first leads to pancreatic enzyme insufficiency (PEI) requiring oral enzyme replacement therapy in 85% of people with CF (Gaskin et al., 1982) and then to CF-related diabetes (CFRD) in around 20% of teenagers and 50% of adults (Moran et al., 2009, Foundation, 2023, Zolin A, 2024, Trust, 2023). CFRD significantly adds to treatment burden and is associated with worse pulmonary function and reduced life expectancy. CFRD is characterised by dysregulated insulin and glucagon secretion (Nielsen et al., 2022, Nielsen et al., 2023). While underlying mechanisms within the pancreas remain unclear, it is increasingly accepted that endocrine dysfunction is at least in part mediated by the severe and progredient pancreatic exocrine pathology (Norris et al., 2019, White et al., 2020).

To date, the studies of the pathological changes within the exocrine and endocrine pancreas in early and late CF have been largely descriptive, with any quantitative analysis focused on endocrine and immune cells (Hart et al., 2018, Bogdani et al., 2017, Löhr et al., 1989). Two distinct pathological patterns have been consistently reported – ‘fibrotic’ and ‘lipoatrophic’ (parenchymal replacement with adipocytes) (Löhr et al., 1989, Bogdani et al., 2017). In parallel with this pancreatic damage, a reorganisation of the islets originally distributed in the acinar parenchyma takes place. This leads to irregular islet clustering although individual islets largely remain identifiable even in end-stage CF (Norris et al., 2019). This suggests that islet reorganisation is secondary to primary CFTR mutation-driven ductal plugging initiating tissue remodelling and ultimately loss of the original compartmentalisation of the pancreas. It is clear that CF pancreatic exocrine pathology progresses incrementally over time but the detailed findings at each stage, the exact sequence of events and putative pathomechanisms have not been systematically characterised.

We have developed a semiquantitative scoring system and AI-driven quantification of pathological findings within the pancreatic exocrine and endocrine compartments to better define and classify the structural changes. The morphometric tools were applied to pancreatic blocks from 29 donors with CF over a wide age range from perinatal death to 27 years’ old in comparison to a control cohort of 58 donors (aged 0 months to 29 years). This enabled mapping of the changes in the pancreatic endocrine and exocrine architecture over the life course of CF. This systematic approach strongly supported a ‘natural history’ of a structural process progressing with age and disease severity from fibrotic changes alone, through a mixed fibrotic / lipoatrophic histological pattern, with culmination in an end-stage lipoatrophic pancreas. Peri- and intra-islet fibrosis in association with activated stellate cells persisted even when the remainder of the gland was entirely replaced by fat, spatially implicating a role for fibrosis and activated stellate cells in diabetes pathogenesis, evidenced by progressive islet remodelling and most severe dysmorphia in association with end-stage exocrine lipoatrophy.

## Methods

### Cohorts

Samples were identified within several pancreatic tissue banks to provide a broad age range in donors with CF and optimally matched control donors. Formalin-Fixed Paraffin-Embedded (FFPE) pancreatic blocks from 19 *post-mortem* donors with CF were provided by the Exeter Archival Diabetes Biobank (EADB). FFPE blocks from 10 *post-mortem* donors with CF were provided from the consultation centre archive of Prof Klöppel (Munich) (Table S1). Ten control deceased organ donor pancreata without CF or diabetes were provided by the MRC Quality in Organ Donation (QUOD) Whole Pancreas Biobank (QUOD PANC) in addition to blocks from 48 *post-mortem* donors without CF or diabetes from EADB (Table S2). *Post-mortem* tissue blocks were collected and biobanked by Dr Alan Foulis (Glasgow, EADB) and Prof Günter Klöppel (Munich). A single block from each donor was analysed except for 6 donors within the Klöppel cohort each providing multiple blocks (Table S1). A subset of donors within the Klöppel cohort were included in a previous publication (Löhr et al., 1989). Deceased organ donor FFPE biopsies were obtained according to quality-assured Standard Operating Procedures from the anterior body region of the pancreas (QUOD designation: P4A; single block from each donor studied)(Kattner et al., 2021). Donor details are given in Table S3 with stains undertaken in each deceased organ donor block summarised in Table S2.

Organs for the MRC QUOD Whole Pancreas Biobank were retrieved after informed and written donor family consent in compliance with the UK Human Tissue Act of 2004 under specific ethical approvals by the UK Human Research Authority (05/MRE09/48 and 16NE0230). The EADB archival tissue bank was approved by the West of Scotland Research Ethics Committee (ref: 20/WS/0074; IRAS project ID: 283620) and the Klöppel cohort archival tissue material by the local ethic committee of the University Hospital ‘rechts der Isar’, Munich, Germany (document number: 281/19 s).

### Staining

Sectioning (at 4 µm thickness) and staining of the QUOD, EADB and Munich consultation archive tissue was performed by NovoPath Laboratories at the Royal Victoria Infirmary, Newcastle upon Tyne. Previously cut sections (4 µm thickness) were provided from the EADB cohort and staining was performed at NovoPath Laboratories. Histological staining for Haematoxylin and Eosin (H&E) and chromogenic staining for chromogranin A (CGA), α-smooth muscle actin (SMA), CD45, CD68, CGA/CD31, insulin/pancreatic polypeptide (PP), and glucagon/somatostatin (Supplementary Table S4) was performed in serial sections using the Ventana Discovery Ultra TM (Roche Diagnostics, Burgless Hill, UK) following NovoPath Laboratory optimised protocols. Sirius Red Fast Green (SRFG) staining was performed following in-house optimised protocol. Stains performed in each donor are itemised in Tables S1 and S2. Stained slides were scanned at 40X magnification at Newcastle Biobank (Leica Aperio AT2 (Leica Biosystems (UK) Ltd)) or in Exeter (Akoya Biosciences PhenoImager Automated Quantitative Pathology Imaging System).

### Histological assessment

#### Semi-quantitative scoring

We devised a semi-quantitative scoring system (based on three grades of severity: mild, moderate and severe) for ductal dilatation, ductal loss, pancreas fibrosis, acinar atrophy, islet remodelling and inflammatory cell infiltration (see Supplementary Table S5). Scoring was undertaken on slides stained with H&E in parallel with SRFG-stained slides for fibrosis evaluation; CGA-stained slides for islet evaluation; and CD45-stained slides to evaluate inflammatory cell infiltration. Descriptors and difficult cases were discussed until a consensus was reached between pathology experts. A key comprising exemplar images for each integer score for each parameter is shown in Supplementary Figures S1-S6.

#### AI-driven quantitative image analysis

Stained sections were analysed with DenseNet AI V2 and Area Quantification modules within the Indica Labs HALO image analysis platform (Version 3.2.1851.354). Quality control was performed, and the following areas were excluded from the analysis: sections of spleen and intestine (in fetal cases), lymph nodes, blurred regions, mucus within large ducts, tears in the tissue and areas with non-specific staining. Individual classifiers were prepared for H&E and SRFG stained sections. The HighPlex BF module was used to enumerate positively stained cells (CD45+) within previous classified regions (endocrine, acinar, fat, fibrosis). The classifiers were trained on donors with a range of CF pathology and three normal pancreata from donors with different ages between <7 days and 3 years. Every example of background, fibrosis, acinar, ducts, islets and fat was annotated manually to ensure accurate training (Supplementary Figure S7). Each classifier was trained to 50,000 iterations and until cross entropy fluctuated between 0.2 and 0.6. Islets were defined as groups of endocrine cells covering an area of ≥1000 µm^2^. Classifiers for islet area were cross-validated in CGA-stained slides. This confirmed excellent correlation between the two methods for quantification of islet area (Supplementary Figure S8). Classifiers for fibrosis area were trained using SRFG-stained slides with all reported collagen density data derived from SRFG-stained slides. The peri-islet region was defined as a ‘halo’ of around 33 µm around the islet. The SRFG-positive area in this region was calculated as a percentage of the total peri-islet area (Supplementary Figure S9). Islet circularity is a measure of the roundness of an islet; it can be calculated when the area and perimeter is known through the formula: C = 4πA/P 2, where C is the circularity, A is the area and P is the perimeter. A perfect circle has C = 1. Quantification of islet hormone positive areas (insulin+, glucagon+, pancreatic polypeptide (PP)+, somatostatin (SST)+) was performed using the HighPlex BF module and presented as a percentage of total of endocrine area (calculated by combining the individual hormone positive areas).

### Statistical analyses

Data are summarised as mean±standard deviation. Comparisons between all CF donors and control donors were undertaken by unpaired Student’s T-test. For comparison of semi-quantitative scores and AI-quantification between tissue blocks of each CF Pattern and control samples, linear mixed effect model was used with CF histological pattern set as fixed effect, donor as random effect and Bonferroni post-hoc testing. Relationships between two variables were assessed using Pearson and Spearman correlation analysis. A p-value of <0.05 was considered statistically significant.

## Results

### Pattern classification of CF pancreas

Each tissue block from the CF donors was classified into one of three patterns: fibrosis without adipocyte replacement (CF Pattern 1: ‘fibrotic’); fibrosis and adipocyte replacement (CF Pattern 2: ‘fibrotic and lipotic’); global adipocyte replacement (CF Pattern 3: ‘lipoatrophic’). Example images for each pattern are shown in Figure 1.

**Figure 1:**
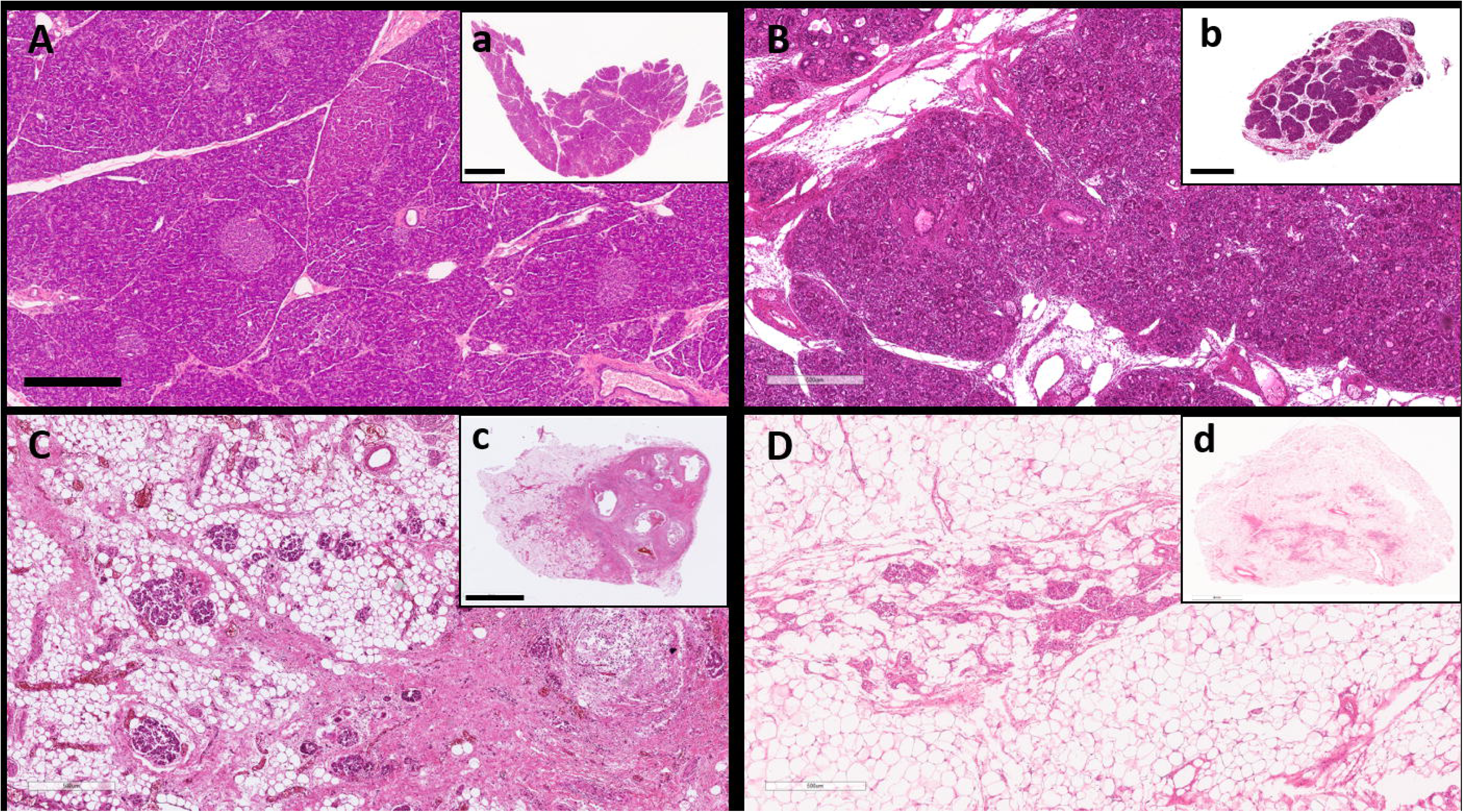
Representative images for control and CF patterns. (A) Control pancreas (Case 53) with normal tissue morphology. (B) CF Pattern 1 with increased collagen deposition but remaining acinar tissue (CF case 9). (C) CF Pattern 2 with extensive acinar atrophy replaced by collagen and adipocytes (CF case 29). (D) CF Pattern 3 with extensive acinar atrophy replaced by fat tissue and clustered islets surrounded by collagen strands (D, CF case 22). Left hand images show higher magnification of each tissue section. Scale bars: A-D 500 µm; a, d 4 mm, b 3 mm, c 5 mm,

In the control pancreas Figure 1A and a and the fibrotic CF Pattern 1 1B and b, the lobularity of the exocrine parenchyma was largely preserved, with only small bands of interlobular fibrosis and single small ducts with mild dilatation and inspissated secretions. The islets were embedded in acinar tissue, with some irregularity of islet shape in comparison to control donors.

In the fibrotic and lipotic pattern (CF Pattern 2, Figure 1C and 1c), there was focal replacement of acinar parenchyma by fibrosis or fatty tissue leading to extensive loss of exocrine parenchyma. Islets remained present within both fibrotic and lipotic areas, though in irregular distribution, shape and size. They were often surrounded and traversed by collagen strands. Ducts and ductules were severely dilated with eosinophilic secretions and mucus in association with extensive peri-ductal fibrosis.

In the lipoatrophic pattern (CF Pattern 3, Figure 1D and 1d), there was total loss of acinar tissue and complete replacement by fat tissue. Islets were still present but aggregated as irregular clusters in the fat tissue intermingled with collagen strands and fat cells. There was extensive duct loss. All remaining ducts were surrounded by fibrosis and mostly small-medium in size (Supplementary Figure S10).

This pattern classification was applied (YA) to each of the 47 samples from the 29 donors with CF. In 20 blocks the tissue changes were classified as Pattern CF1; in 14 blocks as Pattern CF2; and in 13 blocks as Pattern CF3. CF pattern was positively associated with donor age (Spearmans’s rho correlation Coefficient 0.716, p< 0.001) (Supplementary Figure S11).

### Semi-quantitative histological scoring of CF pancreatic changes in comparison to age-matched control donors

Semi-quantitative scoring was undertaken in the 46 CF pancreatic tissue blocks (with the exception of inflammatory cell infiltration scored in the 26 samples stained with CD45) and in samples from 10 age-matched donors without CF, diabetes or pancreatitis. Following unbiased scoring, samples were grouped according to classified Pattern classification (Figure 2).

**Figure 2:**
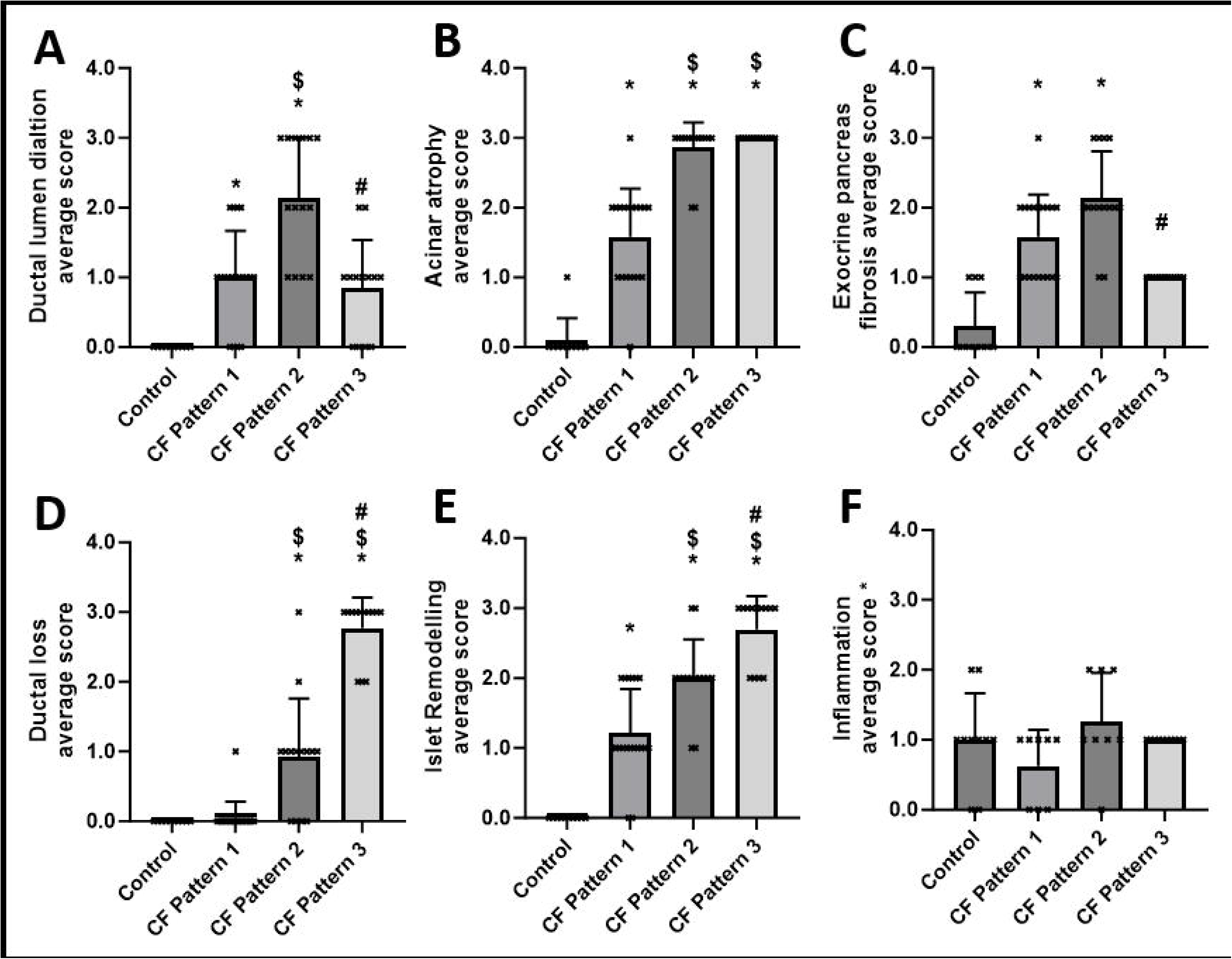
Pattern-based histopathological semi-quantitative analysis to assess control and CF pancreata for different morphological parameters. (A) Ductal lumen dilation, (B) acinar atrophy, (C) exocrine pancreas fibrosis, (D) ductal loss, (E) islet remodelling, and (F) inflammation. Graphs show the mean±SD. Linear mixed effect model (LMEM) was used with CF histological pattern set as fixed effect, donor as random effect and Bonferroni post-hoc test was utilised for statistical analysis. p<0.05 was considered significant. (*) indicates significant difference compared to control, $ indicates significant difference compared to CF Pattern 1, # indicates significant difference compared to CF Pattern 2. A, B, C, D and E: Control n=10 blocks, and CF pancreata n=46 blocks (Pattern 1 n=19 blocks, Pattern 2 n=14 blocks and Pattern 3 n=13 blocks). F control n=10 blocks, and CF pancreata n=26 blocks (Pattern 1 n=8 blocks, Pattern 2 n=8 blocks and Pattern 3 n=10 blocks).

Mild ductal dilatation was evident in CF Pattern 1. Duct dilatation was ubiquitous in Pattern 2 with residual ducts in Pattern 3 having relatively little dilatation. The exocrine Pancreatic Fibrosis Score closely paralleled the Ductal Dilatation Score, being significantly greater than in control donors without CF in Pattern 2 and peaking in Pattern 2. This appeared to be followed by duct loss with no difference in Duct Loss Score from control donors in Pattern 1, ‘focal’ areas of loss in Pattern 2 and ‘severe’ loss in Pattern 3. Acinar atrophy was already marked in Pattern 1 and was virtually complete in both Pattern 2 and 3.

The islet remodelling score (Supplementary Table S5) reflected disease progression from fibrosis alone, through a combined fibrotic / lipotic phenotype, to global lipoatrophy evidenced by incrementally greater islet aggregation associated with increasing variability in size and shape. Peri-islet fibrosis was notable in CF Pattern 2 and 3 in addition to intra-islet fibrosis, with 5.8±4.7% (Pattern 2) and 8.6±6.8% (Pattern 3) of islets (on manual quantification) affected by significant fibrosis extending beyond the intra-islet perivascular region (Supplementary Figure S12, S13).

Semi-quantitative scores for all features were significantly higher in CF compared to control with the exception of inflammatory cell infiltration. ‘Mild inflammatory cell infiltration’ was present in control donors and in CF without presence of ‘severe inflammation’ in any of the three histological patterns (Figure 2F).

### AI analysis of pancreatic exocrine compartment

Quantification of stained sections from 25 CF donors in comparison to 58 control donors was undertaken following training of the Indica Halo AI deep learning classification tool. Statistical comparison of all CF donors versus age-matched control donors was undertaken and data were plotted according to donor age. Comparison of all blocks classified into each of the three patterns versus control donors was performed.

Approximately 80% of pancreatic area comprised acinar tissue in control donors compared with mean area of <20% in donors with CF (p<0.0001) (Figure 3A1). Perinatally, acinar area extent was similar to that of control donors (Figure 3A2), in keeping with significant stromal tissue area during normal as well as CF embryonic development. Acinar area was, however, significantly reduced in some very young donors with CF. Acinar tissue was completely absent in CF donors by 8 years of age. A significant reduction in acinar area was seen in sections with CF Pattern 1, with virtually no remaining acinar tissue in sections with Pattern 2 or 3 (Figure 3A3).

**Figure 3:**
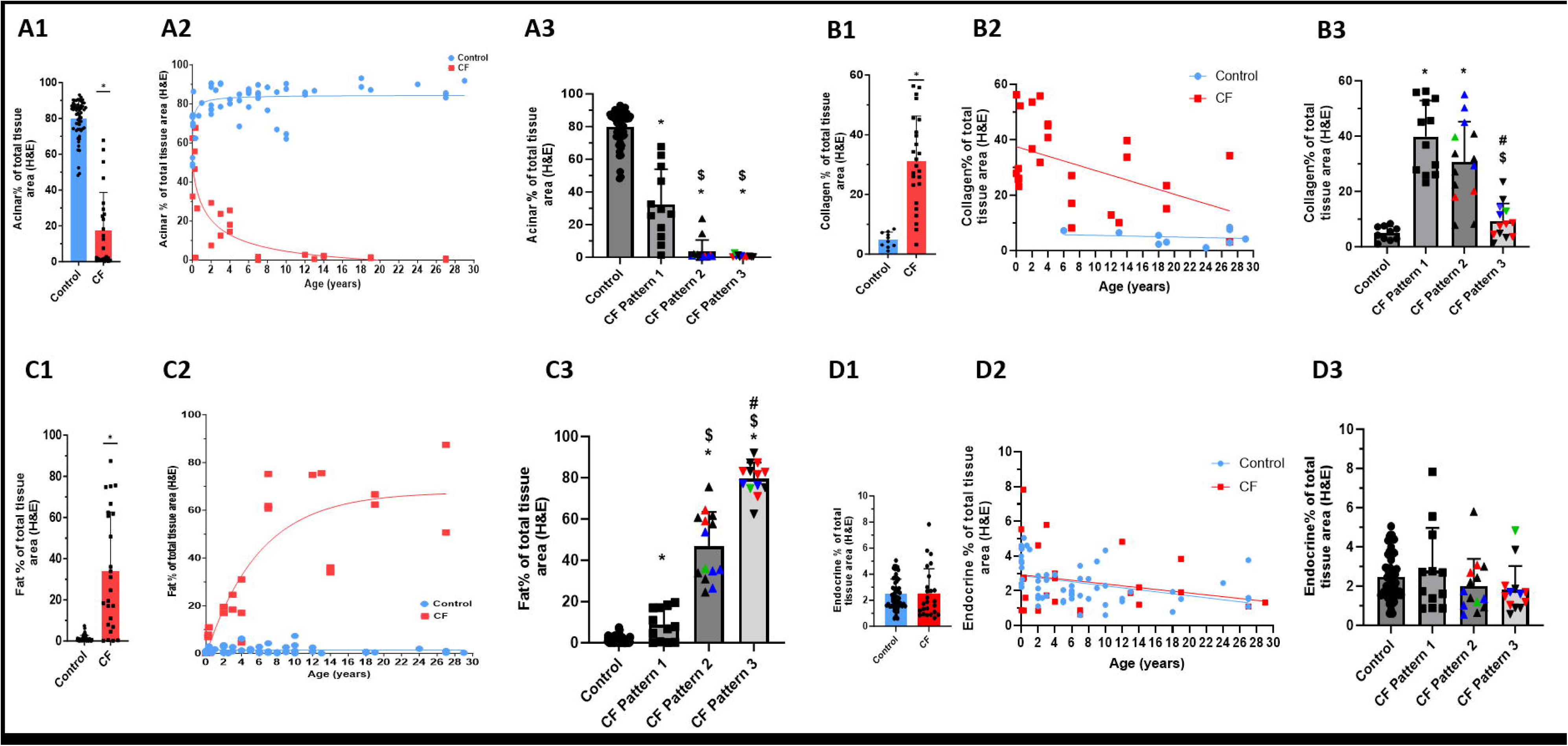
AI-based analysis of pancreatic histopathological parameters: acinar PA (A1-A3); collagen PA (B1-B3), adipocyte PA (C1-C3), and endocrine PA (D1-D3). All control (blue) and CF (red) donors were compared by unpaired Student’s T-test (A1, B1, C1, D1). Donor-based analysis vs age in control (blue circles) and CF (red squares) donors (A2, B2, C2, D2). Block-based analysis of control vs CF Pattern 1, 2, and 3 (A3, B3, C3, D3). Linear mixed effect model (LMEM) was used with histological CF pattern set as fixed effect, donor as random effect and Bonferroni post hoc test was utilised for statistical analysis. Graphs show mean±SD. p<0.05 was considered statistically significant. (*) indicates significant difference compared to control, ($) significant difference compared to CF Pattern 1, # indicates significant difference compared to CF Pattern 2. A3, B3, C3, D3 Data points coloured red represent Case 22 and points coloured blue represent Case 29, both donors have multiple blocks of different patterns. Points coloured green represent CF Cases 21 and 24 with diagnosed CFRD. A1, A2, C1, C2, D1, D2: control n=58 donors, CF n=25 donors. B1, B2: control n=10 donors, CF n=25 donors. A3, C3, D3: control n=58 blocks, CF Pattern 1 n=12 blocks, CF Pattern 2 n=14 blocks, CF Pattern 3 n=13 blocks. B3: control n=10 blocks, CF Pattern 1 n=12 blocks, CF Pattern 2 n=14 blocks, CF Pattern 3 n=13 blocks.

Collagen area was significantly higher in donors with CF comprising >30% of overall area versus 5% in control donors (p<0.0001) (Figure 3B1). Percentage collagen area decreased with CF donor age (B2) in parallel with increasing (Pattern 2) and ultimately global (Pattern 3) adipocyte replacement. Overall collagen area in Pattern 3 was no longer significantly greater than in controls (Pattern) (B3).

Mean adipocyte area was 34% in CF donor pancreata compared with only 1% in control donors (p<0.0001) (Figure 3C1). Adipocyte areas in CF donors were greater than controls even in youngest donors increasingly rapidly with age (C2) and from CF Pattern 1 through Pattern 2 and ultimately Pattern 3 (C3). The percentage of adipose tissue in control pancreata remained very low regardless of age.

AI analysis for ductal area could not be meaningfully completed because the classifier could not accurately distinguish between blood vessels and pancreatic ducts.

### AI analysis of pancreatic endocrine compartment

The proportional endocrine area per section was comparable between control (2.5±1.2%) and CF (2.5±1.9%) cohorts (p=0.873) and remained relatively stable with age in both groups (Figure 3D2). There was no statistically significant differences between the different CF patterns (Figure 3D2). These findings are in keeping with more rapid expansion of exocrine in comparison to endocrine area in the growing pancreas (in the absence of disease) without significant loss of overall islet volume in progressive CF pathology.

Islet density (calculated as number/mm^2^ total tissue area) was also comparable between control (4.6±2.7 islets/mm^2^) and CF (4.6±4.0 islets/mm^2^) pancreata overall (Figure 4A1). This decreased markedly with increasing age in both groups (Figure 4A2). Islet density in sections with CF Pattern 2 and 3 was significantly lower than in CF Pattern 1 (Figure 4A3).

The size of individual islets (determined by islet diameter (Figure 4B) and area (Figure 4C)) was not significantly different between CF and control donors but both increased linearly with donor age. Islets were significantly larger in CF Patterns 2 and 3 compared to CF pattern 1 (Figure 4B3 and 4C3). This may reflect islet growth in both groups or expansion of islet diameter and area by interposing fibrous tissue between endocrine cells in advanced CF. The ability of AI to successfully differentiate individual islets within clusters was evidenced by highest median diameters of approximately 150 µm. Median islet circularity was lower in CF versus control donors (Figure 4D1) without any significant changes with donor age (Figure 4D2). AI-determined circularity was comparable between all CF Patterns (Figure 4D3).

**Figure 4:**
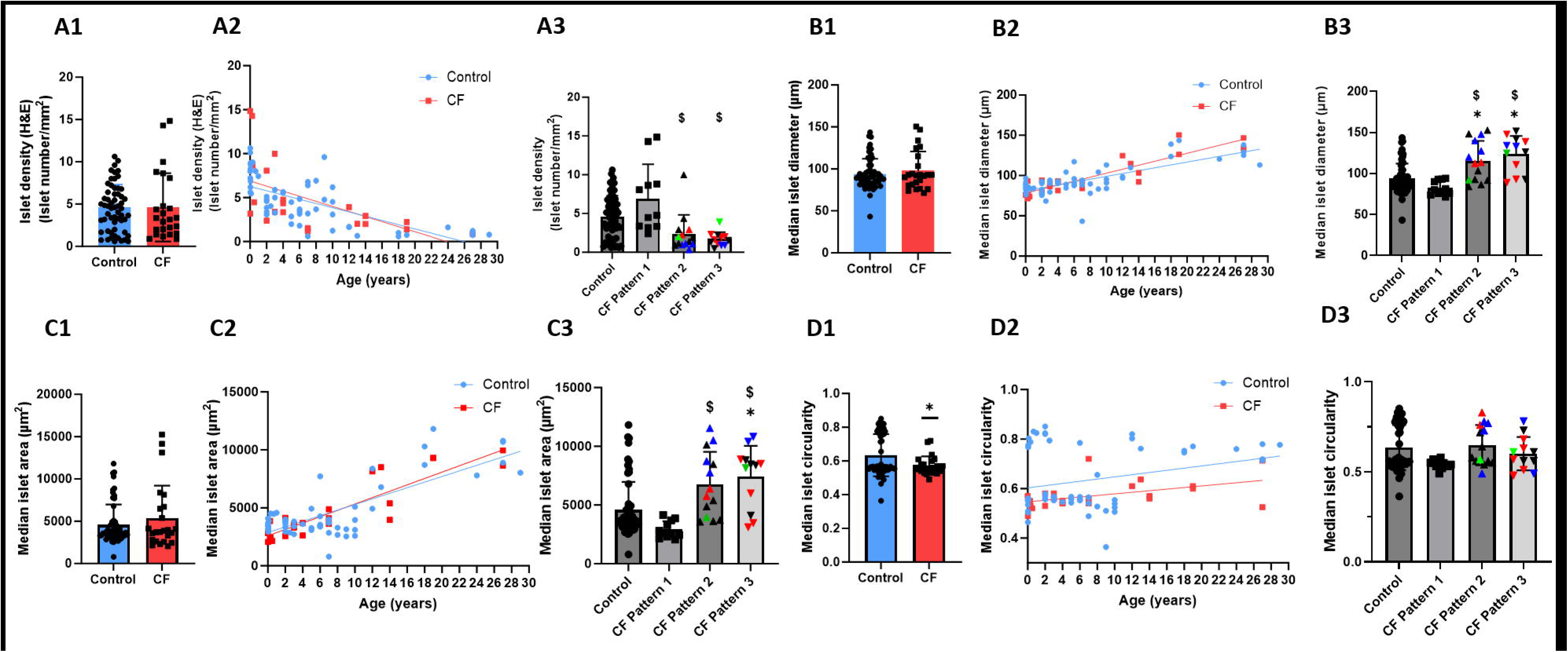
AI based analysis of islet density (A1-A3), islet diameter (B1-B3), islet area (C1-C3), islet circularity (D1-D3). All control (blue) and CF (red) donors were compared by unpaired Student’s T-test (A1, B1, C1, D1). Donor-based analysis vs age in control (blue circles) and CF (red squares) donors (A2, B2, C2, D2). Block-based analysis of control vs CF Pattern 1, 2, and 3 (A3, B3, C3, D3). Linear mixed effect model (LMEM) was used with Pancreatic Pattern set as fixed effect, donor as random effect and Bonferroni post hoc test was utilised for statistical analysis. Graphs show mean±SD. p<0.05 was considered statistically significant (*) indicates significant difference compared to control, ($) significant difference compared to CF Pattern 1. A3, B3, C3, D3. Data points coloured red represent Case 22 and points coloured blue represent Case 29, both donors have multiple blocks of different patterns. Points coloured green represent CF Cases 21 and 24 with diagnosed CFRD. A1, A2, B1, B2, C1, C2, D1, D2: control n=10 donors, CF n=24 donors. A3, B3, C3, D3: control n=58 blocks, CF Pattern 1 n=11 blocks, CF Pattern 2 n=14 blocks, CF Pattern 3 n=12 blocks.

### Changes to endocrine tissue composition in the CF pancreas

To investigate proportions of individual endocrine cell phenotypes within islets, sections from a subset of 22 CF blocks and 10 control blocks were stained by double-immunohistochemistry for both insulin, to identify β-cells (brown) and PP, to identify PP cells (red), with the next serial section stained for both glucagon, to identify α-cells (brown) and somatostatin, to identify δ-cells (red) (Figure 5A). The percentage area for each hormone as a proportion of the overall immunostained endocrine area (four stains combined), termed proportional area (PA) was calculated (Figure 5, Figure S14). Donor-level analysis demonstrated a significant decrease in insulin PA in CF donors (49±17%) compared to control donors (83±6%) (p<0.0001) (Figure 5B1). In parallel, the glucagon and somatostatin PA were significantly increased in CF donors (Figure 5C1, Figure 5D1). Glucagon PA significantly increased from 9±2% in control to 29±11% in CF donors (p<0.0001) while somatostatin PA increased from 7±4% in control to 24±9% in CF donors (p<0.0001). Pancreatic polypeptide PA was comparable between CF and control donors (Figure 5E1).

**Figure 5:**
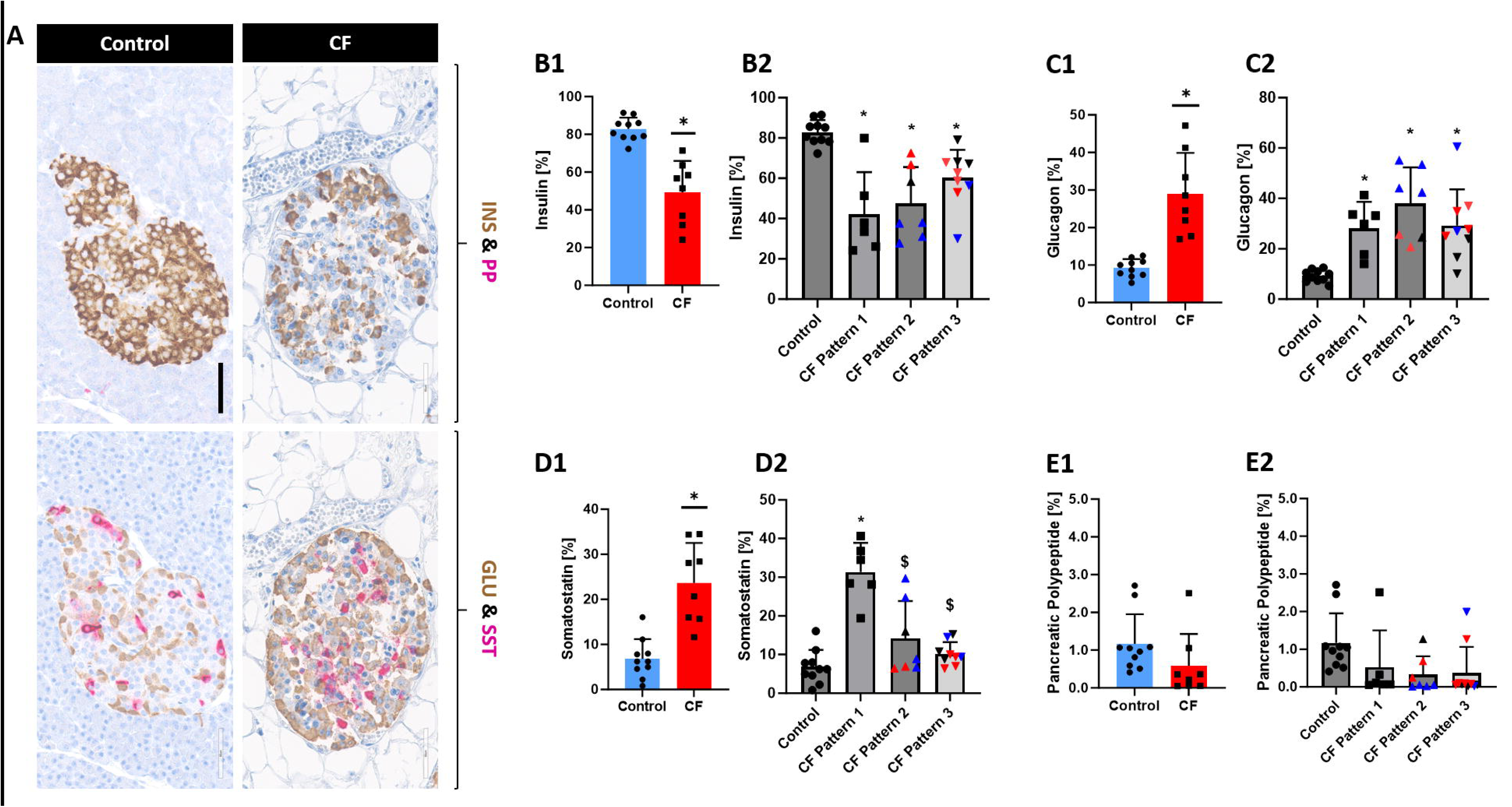
AI analysis of islet hormone composition in CF and control pancreata. (A) Dual IHC staining for insulin (brown) / PP (pink) and glucagon (brown) / somatostatin (pink) in a control (Case 53) and CF pancreas (Case 29). Scale bar 60 µm. Donor-based AI analysis of insulin (B1), glucagon (C1), somatostatin (D1), and PP (E1) PA in control (blue) and CF (red) pancreata. p<0.05 was considered significant. (*) indicates significant difference compared to control. Control n=10 donors, CF n=8 donors. Block based AI quantification of insulin (B2), glucagon (C2), somatostatin (D2), and PP (E2) PA. Control n=10 blocks, CF n=22 blocks (Pattern 1 n=6, Pattern 2 n=7 and Pattern 3 n=9). Donor-based comparison was carried out using unpaired Student’s T test and block-based comparisons were conducted using linear mixed effect model (LMEM) with histological CF pattern set as fixed effect, donor as random effect and Bonferroni post hoc test. Graphs show the mean±SD. p<0.05 was considered significant. (*) indicates significant difference compared to control. Data points coloured red represent Case 22 and blue coloured points represent Case 29. Both donors have multiple blocks with different CF Patterns.

Plotting PA for each endocrine hormone versus donor age demonstrated that decreased β-cell and increased α-cell PA were noted from birth in CF and this pattern was maintained with increasing age (Supplementary Figure S14A and B). Somatostatin PA was highest perinatally and decreased with age (Supplementary Figure S14C).

Reduction in insulin PA and increase in glucagon PA versus control blocks remained constant in all CF patterns without any evidence of further proportional β-cell loss with exocrine disease progression (Figure 5B2 and C2). Somatostatin PA was significantly higher than control blocks in CF pattern 1 but decreased significantly in association with CF patterns 2 and 3 (Figure 5D2). No significant differences in pancreatic polypeptide PA were seen between any CF pattern and control blocks (Figure 5E2, Supplementary Figure 14D).

### Increased fibrosis in the islet microenvironment in CF

Peri- and intra-islet collagen deposition was significantly higher in CF cases in comparison to control donors (Figure 6). Despite the small numbers of donors studied, persistence of intra-islet fibrosis even in the oldest donors was seen (Figure 6C2, C3). Although no peri-islet fibrosis data were available for donors in CF Pattern 1, block-based analysis revealed that both peri-islet (Figure B3) and intra-islet (Figure C3) collagen density were highest in CF Pattern 2 persisting in Pattern 3.

**Figure 6:**
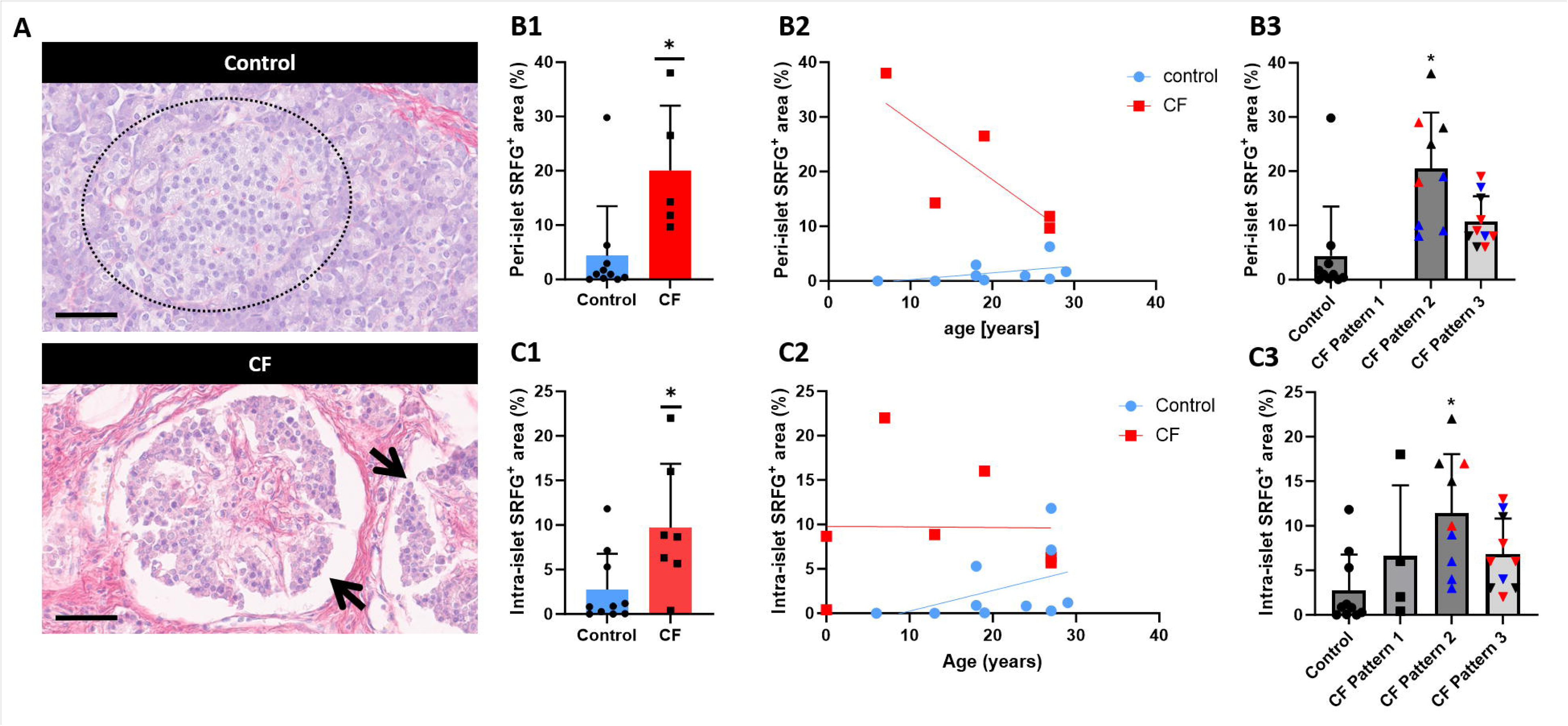
Collagen in the islet microenvironment in CF and control pancreata. (A) Representative images of SRFG staining of a control (Case 50) and CF (Case 22) pancreas. Dashed circle indicates islet in control tissue, black arrows indicate islets in CF tissue. Scale bar 60 µm. Donor-based AI analysis of the peri-islet (B1) and intra-islet (C1) proportion of collagen in control (blue) and CF (red) pancreata. B1: Control n=10 donors, CF n=5 donors. C1: Control n=10 donors, CF n=7 donors. Block-based AI analysis of peri-islet (B2) and intra-islet (C2) proportion of collagen. B2: Control n=10 blocks, CF n=19 blocks (Pattern 1 n=0, Pattern 2 n=9 and Pattern 3 n=10). C2: Control n=10 blocks, CF n=23 blocks (Pattern 1 n=4, Pattern 2 n=9 and Pattern 3 n=10). Donor-based statistical analyses were carried out using unpaired Student’s T test and block-based analyses were conducted using linear mixed effect model (LMEM) with CF histological pattern set as fixed effect, donor as random effect and Bonferroni post-hoc test. Graphs show the mean±SD. p<0.05 was considered significant. (*) indicates significant difference compared to control. Data points coloured red represent Case 22 and blue coloured points represent Case 29. Both donors have multiple blocks with different CF Patterns.

#### Cellular component to fibrosis

Inflammatory cells in pancreatic tissue of both CF and non-CF donors were detected by AI-assisted analysis following CD45 immunohistochemical staining (Supplementary Figure 15). The overall leucocyte number was significantly lower in CF donors with 43±40 cells/mm^2^ (n=5) compared to control donors with 282±123 cells/mm^2^ (n=10) (p<0.01) (Supplementary Figure 15A). Although numbers were low, immune cell counts appeared relatively consistent regardless of age in control donors and did not increase with age in CF pancreata (Supplementary Figure 15B). Numbers were also comparable between CF patterns 2 and 3 with no data available for Pattern 1 (Supplementary Figure 15C).

Leucocytes, mainly lymphocytes and histiocytes, could be identified in all slides examined (with the exception of a single donor with CF (Case No. 3)). Numbers were higher around vascular structures with an example in a control donor shown in Supplementary Figure 15D1 (dashed oval). In many CF cases, foci of inflammation were observed (Supplementary Figure 15D2). In some donors with CF, there was evidence of increased leucocyte density around ducts and islets (Supplementary Figure 15D3).

In addition to the systematic quantitative analysis of pancreatic fibrosis and inflammatory cell distribution, a subset of CF and control pancreata were stained for CD68 and α-SMA to further investigate cellular associations with fibrosis.

Seventeen CF pancreatic blocks were stained for CD68 to assess the distribution of macrophages. These where found scattered in the stroma, within inflammatory cell foci and with a peri- and intra-islet distribution in some sections (Supplementary Figure S16).

Alpha-SMA staining was used as a marker for myofibroblastic cells which correspond to activated pancreatic stellate cells (Omary et al., 2007) in CF (n=31) and non-CF pancreata (n=9) (Figure 7). Positive cells were observed in 9 control pancreata around ducts (Figure 7A) and around islets (Figure 7B, arrow). In the 31 CF pancreata, we observed an increase in the number and density of activated pancreatic stellate cells in areas of fibrosis surrounding ducts (Figure 7C) and islets (Figure 7D). Cellular morphology and topography in addition to CD31 immunostaining was used to differentiate stellate from endothelial cells and other perivascular cells.

**Figure 7:**
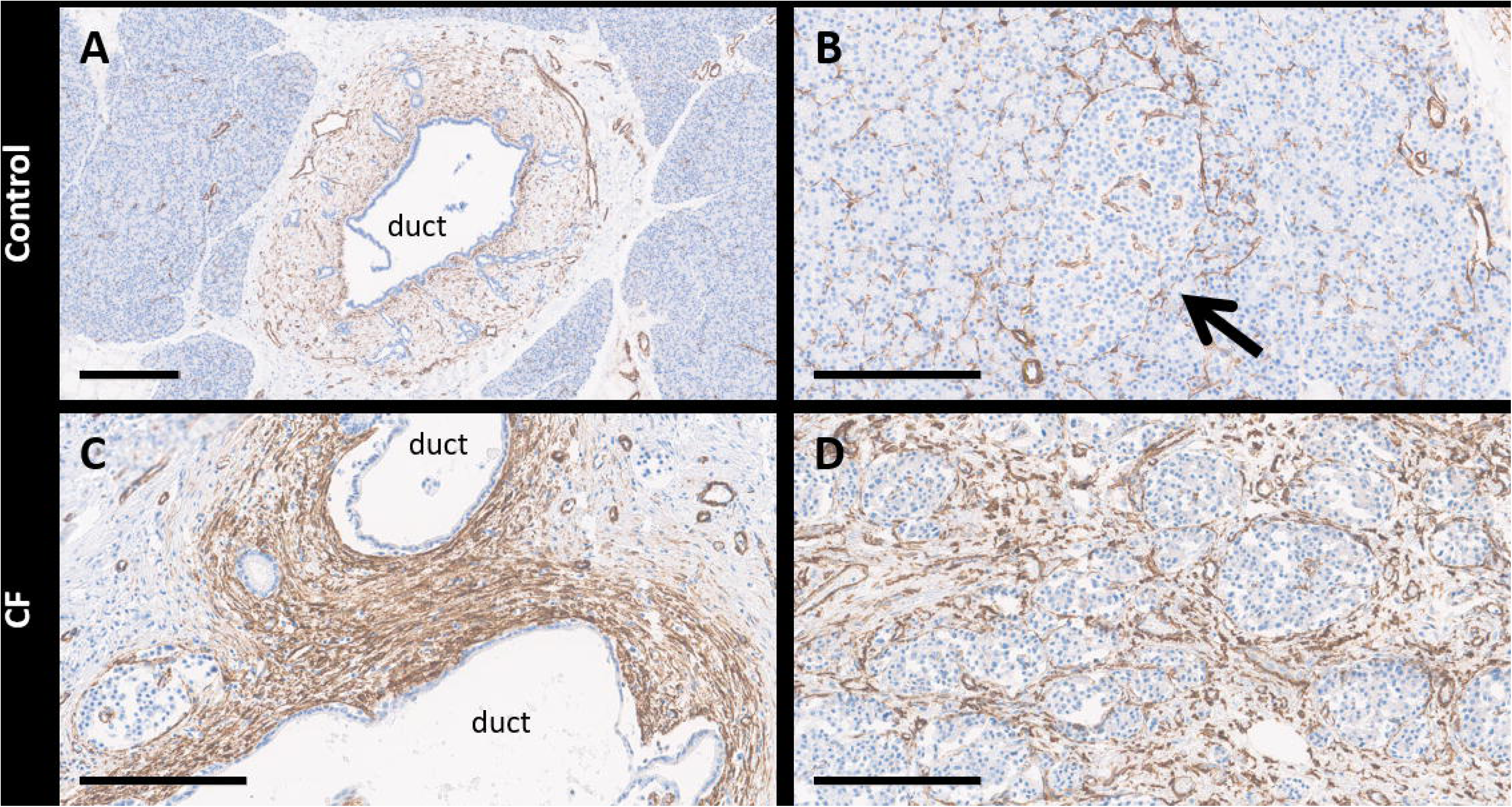
Distribution of stellate cells in CF and control pancreata. Immunostaining for α-SMA (brown) in control (A, B) and CF (C, D) pancreata indicating pancreatic stellate cells. (A, C) α-SMA in the periductal stroma. (B, D) α-SMA in the islet microenvironment. Black arrow indicates islet in panel B. A: Donor number 31, B: Donor number 51, C: Donor number 22, D: Donor number 27. Scale bars: A – 300 µm; B,C,D – 200 µm.

## Discussion

Pancreatic pathology of CF has been described qualitatively in several studies (Löhr et al., 1989, Couce et al., 1996, Abdul-Karim et al., 1986, Cory et al., 2018). However, there is yet no detailed quantitative approach which allows a refined but standardised assessment of the pathological changes within exocrine and endocrine compartments in the CF pancreas. As a result, no clear timeline for the changes could be determined. To address this, we undertook a systematic approach to classify morphological patterns, developing both a semi-quantitative histological score and a complementary AI-based quantification of histological features in CF pancreata. We applied this to a CF cohort with a wide age range in parallel to age-matched controls. We compared CF with control donors, plotted data according to donor age and undertook comparative analysis according to morphological patterns of remodelling.

Development and application of a light microscopy scoring system separately assigning integer scores to specific parameters and training AI software to classify and quantify exocrine and endocrine compartments in CF and control tissue sections has provided a robust approach to systematic unbiased analysis of pancreatic pathology. While semiquantitative scores for fibrosis (Bogdani et al., 2017) and other exocrine changes including adipocyte replacement (Iannucci et al., 1984) (Soejima and Landing, 1986) have previously been utilised to report CF pancreatic pathology, automated AI quantification of exocrine parameters has not been reported. AI evaluation of acinar atrophy and fibrosis in CF samples with fibrosis alone (CF Pattern 1), fibrosis and adipocyte replacement (CF Pattern 2) and lipoatrophy (CF Pattern 3) mirrored semi-quantitative analysis in the current and previous studies (Iannucci et al., 1984, Soejima and Landing, 1986, Bogdani et al., 2017) but enabled for the first time proportional quantification of the progressive changes in each section of the CF pancreata.

The two patterns of exocrine pancreatic pathology in adults with CF - ‘fibrotic’ and ‘lipoatrophic’ are well-recognised in addition to an association between CFRD and extensive fat replacement (Löhr et al., 1989, Iannucci et al., 1984, Bogdani et al., 2017). Whether fibrosis is a precursor to ‘inevitable’ fat replacement has remained unclear and debated. However, the rapid increase in AI-quantified adipocyte PA with increasing age culminating in virtual complete replacement of the exocrine pancreas by fat tissue demonstrated in the current study provides convincing evidence for inexorably progressive fat replacement in the CF pancreas during prolonged complete duct blockage by viscous mucin. This is in keeping with imaging studies demonstrating replacement of the gland almost entirely by fat in all adults with CF and pancreatic exocrine enzyme insufficiency (Daneman et al., 1983, Soyer et al., 1999). In a pilot study in Copenhagen (Niessen B, Faurholt-Jepsen D unpublished data) in nine people with CF and pancreatic exocrine enzyme insufficiency aged >18 years’ old, pancreatic fat percentage measured by magnetic resonance imaging was >75% regardless of glucose tolerance - ranging from normal to CFRD. In a further study an association between pancreatic fat replacement and degree of exocrine but not endocrine function was shown (Soyer et al., 1999). Moreover, Shwachman’s syndrome, a hereditary disorder associated with replacement of pancreatic parenchyma with adipocytes without fibrosis is associated with exocrine enzyme deficiency but not diabetes (Tirkes et al., 2019, Löhr et al., 1989). While the aetiological drivers of fat replacement in CF remain unknown, we have detected upregulation of genes associated with fatty acid metabolism and adipogenesis in Nanostring analysis of pancreata within the current cohort with CF Pattern 2 and 3 (unpublished data). Upregulation of adipogenic genes within the pancreas has also been described in a ferret model of CF (Yi et al., 2016).

Our analysis robustly evidences a natural history progressing from ductal dilation and expanded interlobular stroma progressing from birth, followed by rapid loss of acinar PA. Peri-ductal fibrosis is followed by sub-total ductal loss. Ductal and acinar volume in addition to confluent fibrosis becomes increasingly replaced by adipocytes.

Although previous publications have emphasised the relative normality of exocrine architecture in very young individuals with CF, our analysis provides evidence for a natural history of CF changes in the pancreas starting from and likely preceding birth. In line with the findings of Bogdani et al., (Bogdani et al., 2017) in CF infants under 1 year of age, we found, as earliest significant changes in the CF pancreas in comparsion to control donors, distinctly dilated ducts in addition to widening of the interlobular area with deposition of small collagen bands. These findings suggest that the remodelling process in the CF pancreas has its origin in the duct cell, an assumption supported by strong CFTR expression in the human pancreatic duct cells (Di Fulvio et al., 2020, White et al., 2020, Marino et al., 1991).

Together with duct changes, a relative decrease in acinar volume has been reported even at birth (Imrie et al., 1979) with increasing acinar loss with age in very young CF donors (Bogdani et al., 2017). We quantified the rapid rate of acinar cell loss over a wide age range and demonstrated near-complete acinar atrophy by the age of 7 years. This has been confirmed clinically by early onset of pancreatic exocrine insufficiency in the majority of people with CF (Gaskin et al., 1982, Singh and Schwarzenberg, 2017, Ahmed et al., 2003, Zolin A, 2024, Walkowiak et al., 2005).

Pancreatic fibrosis is a central finding in CF (Iannucci et al., 1984, Soejima and Landing, 1986, Löhr et al., 1989, Couce et al., 1996, Bogdani et al., 2017). In our study the fibrosis started in the periductal area, with its severity mirroring the degree of ductal dilatation. Exocrine fibrosis was greatest in CF Pattern 1 and decreased in older donors as previously described (Iannucci et al., 1984, Löhr et al., 1989) with involution paralleling ductal loss. Residual small ducts in end stage lipoatrophic CF were in all cases surrounded by persisting fibrosis.

Peri- and intra-islet fibrosis were evident in the current analysis peaking with CF Pattern 2. This has been observed previously but not quantified (Iannucci et al., 1984, Abdul-Karim et al., 1986, Löhr et al., 1989, Couce et al., 1996, Bogdani et al., 2017, Hart et al., 2018). Through our systematic approach we were able to confirm persistent presence of peri- and intra-islet fibrosis in the Pattern 3 lipoatrophic gland, will collagen making up a significant proportion of overall islet area.

We have confirmed a cellular component to the initial peri-ductal and later peri- and intra-islet fibrosis in CF through presence of α-SMA-positive activated PSCs. In addition to perivascular leukocytes in CF and presence of infiltrating inflammatory cell foci comprising predominantly lymphocytes and histiocytes, peri-ductal and peri-islet leukocytes including macrophages were seen.

Increased numbers of immune cells including macrophages around and within islets have previously been reported in addition to β-cell expression of the inflammatory cytokine IL1-β (Bogdani et al., 2017, Hart et al., 2018, Hull et al., 2018).

The number of pancreatic immune cells in CF donors quantified by AI in the current study was comparable to a previous report (Bogdani et al., 2017). In our study, immune cell count was surprisingly high in the age-matched controls without CF. None had evidence of other overt pathology but all were characterised by high exposure to inflammatory stressors around the time of death and donation (Supplementary Table S4). Cause of death was hypoxic brain injury in 60% of cases with cardiorespiratory arrest in 70% and elevation of the acute phase reactant C-reactive peptide in 80%. These factors may provide at least partial explanation for the relatively high numbers of inflammatory cells in these donors.

Activated PSCs in association with macrophages are increasingly believed to play a critical role in pancreatic fibrogenesis and tissue remodelling in chronic pancreatitis and animal models (Haber et al., 1999, Omary et al., 2007, Baron et al., 2016, Casini et al., 2000, Bachem et al., 1998, Zang et al., 2015, Glaubitz et al., 2023). Peri-ductal activated PSCs have been identified in the ferret CF model but not in age-matched wild type controls and it has been hypothesised that these are key to pancreatic exocrine destruction and adipogenesis (Rotti et al., 2018).

In a comparison between 25 individuals with CF representing a wide spectrum of age and 58 age-matched controls, AI quantification demonstrated no significant differences between percentage endocrine area, islet density, median islet diameter or area. Islet density appeared to reduce and median islet size increase with increasing age in both CF and normal donors. CF Patterns 2 and 3 associated with older age were also associated with reduced islet density and increased size in comparison to Pattern 1. Whether this represents different processes remains unclear: increasing islet volume being ‘outstripped’ by increasing pancreatic exocrine volume leading to reduced density but increased pancreatic endocrine mass during normal growth; and true loss of islet mass reflected by decreasing density within a diseased non-growing pancreas in CF with increasing size of individual islets reflecting an increase in interposing intra-islet fibrosis rather than increasing endocrine mass. It is notable that assessment of islet area in CF by the primary method used in our study of AI classification based on H&E staining (capturing overall islet area) yielded higher areas than measurement of true endocrine cell area using chromogranin A staining despite very good overall correlation between the two methods (Supplementary Figure S9). The absolute values and slopes with age were remarkably closely aligned between CF and non-CF, however (Figure 3D). Moreover, overall percentage endocrine area was not significantly lower with increasing age or progression from Pattern 1 to Pattern 3 in CF, as previously reported in quantitative studies (Löhr et al., 1989, Cory et al., 2018, Bogdani et al., 2017). However, a decrease in endocrine mass was observed in CFRD compared to CF in two studies with smaller cohorts (7-8 cases of CFRD) (Iannucci et al., 1984, Bogdani et al., 2017). Through our systematic well-controlled unbiased AI analysis, we can conclude that although some loss of overall islet mass cannot be ruled out, this is at worst modest.

Mean islet circularity determined by AI was lower in CF than in normal controls. Our semi-quantitative scoring system revealed progressive remodelling of the pancreatic endocrine compartment with peri-islet fibrosis in Pattern 1; islet aggregation within complexes surrounded by fibrosis and activated PSCs in Pattern 2; and islets with variable size and shape including a proportion with extensive intra-islet fibrosis in Pattern 3. Even in the presence of global fat replacement and gland atrophy, all islet clusters were associated with surrounding collagen strands and residual fibrosis. Although the presence of peri-islet fibrosis in progressive CF has previously been reported (Iannucci et al., 1984, Abdul-Karim et al., 1986, Löhr et al., 1989, Couce et al., 1996, Bogdani et al., 2017), some groups have concluded that the fat replacement which is greatest in those with CFRD is a primary driver (Bogdani et al., 2017) although in our study adipocyte replacement appeared to be virtually complete in relatively young donors, likely preceding diabetes development. We hypothesise that peri- and intra-islet fibrosis are playing a key role in pancreatic endocrine dysfunction and ultimately in progression to diabetes and that this process may be mediated by activated pancreatic stellate cells. We propose that activated pancreatic stellate cells are orchestrating reorganisation of the islet niche interrupting normal endocrine cell-to-cell communication critical for normal function and potentially also normal blood flow through vascular ‘strangulation’ (Löhr et al., 1989). In addition to fibrogenesis, PSC secretome may directly impair proliferation, viability and function of endocrine cells (Zang et al., 2015, Zha et al., 2014, Brozzi et al., 2015, Oleson et al., 2015).

Within islets, insulin PA was reduced by almost 50% in Pattern 1 CF in comparison to control donors without CF. This was the case even in donors dying peri-natally and at 4 months of age, in agreement with a previous study investigating young CF donors aged between 5 days and 4 years (Bogdani et al., 2017). This reduced proportional β-cell area remained relatively constant with CF donor age and in Pattern 2 / 3. The absence of further β-cell loss during development of abnormal blood glucose levels and even after progression to CFRD is reflected in all other published literature relating to human donors (Abdul-Karim et al., 1986, Bogdani et al., 2017, Cory et al., 2018, Couce et al., 1996, Hart et al., 2018, Löhr et al., 1989). Decreased insulin PA was mirrored by a three-fold increase in glucagon PA and this again remained relatively stable with age and exocrine pancreatic disease progression in keeping with the majority of previous publications (Bogdani et al., 2017, Hart et al., 2018, Hull et al., 2018, Löhr et al., 1989). Somatostatin PA was also three-fold higher in Pattern 1 CF donors compared with non-CF controls as previously reported (Bogdani et al., 2017, Hull et al., 2018, Löhr et al., 1989) but decreased with age and CF disease progression in contrast to insulin and glucagon PA.

Strengths of our study include the relatively large number of donors and age-matched controls obtained by pooling several tissue repositories over a wide age range and applying a systematic approach including semi-quantitative analysis informed by expert pathologists and then applied consistently across all tissue sections to minimise bias, in parallel with unbiased AI image analysis. This approach has potential for application beyond CF for robust scoring of both exocrine and endocrine compartments in other pancreatic diseases including chronic pancreatitis (Esposito et al., 2020). Weaknesses include autolytic changes within *postmortem* tissue affecting staining quality. Further, gross disruption of pancreatic parenchyma in advanced CF impacted the accuracy of AI classification, leading to exclusion of all quantitative pancreatic duct data. In addition, limited donor data were available for the historic CF cases, with *antemortem* diabetes status and CF genotype unknown.

Through cross-sectional systematic analysis in a relatively large cohort of donors with CF and age-matched controls pooled from several international tissue banks, we have obtained detailed findings from the early and advanced phases of disease allowing us to construct a morpho-functional timeline of the natural history of pancreatic CF. From this we have made three conclusions regarding the potential drivers of β-cell dysfunction and CF-related diabetes:

1. Although it appears that reduced β-cell mass is established at birth, progression through decreasing β-cell function to diabetes is not associated with any further proportional β-cell loss (Figure 8A).
2. Fat replacement increases with age with extent associated with acinar / ductal loss and thus loss of exocrine function through pancreatic exocrine enzyme deficiency but not loss of endocrine function and progression to diabetes (Figure 8B)
3. Pancreatic fibrosis follows a biphasic pattern with predominantly peri-ductal fibrosis during development of pancreatic exocrine insufficiency. This is followed by peri-islet fibrosis enriched with activated stellate cells and the presence of macrophages associated with islet clustering and progressive disorganisation. This process culminates in extensive intra-islet collagen deposition potentially interrupting normal endocrine cell-to-cell communication and restricting islet blood flow. We propose this cellular fibrotic process in and around the islet compartment in later stage pancreatic CF as a primary driver of progression to CFRD (Figure 8C).

**Figure 8:**
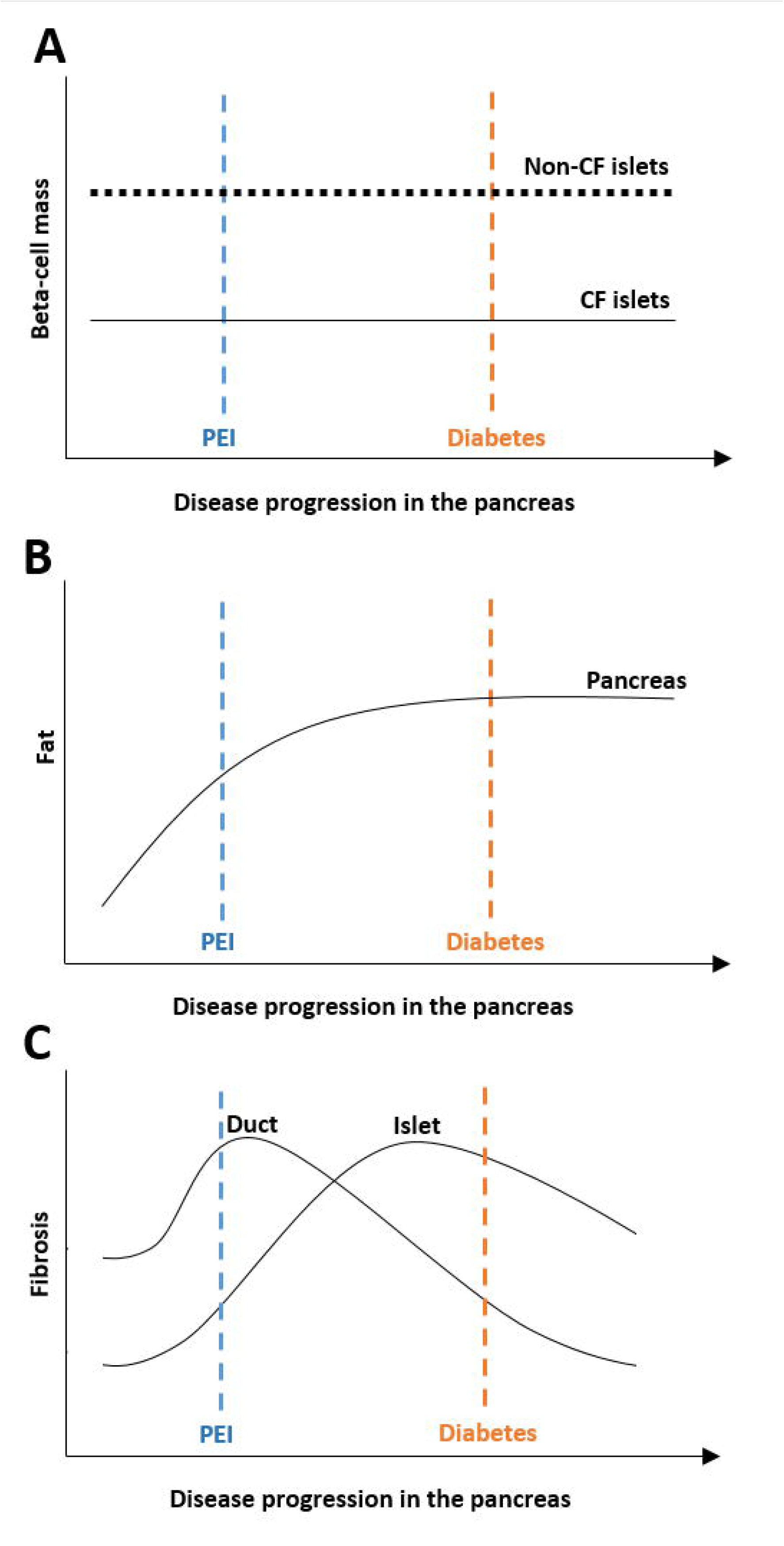
Schematic of β-cell mass (A); adipocyte replacement (B); and fibrosis in the microenvironments of ducts and islets (C) in relation to functional disease progression of CF within the pancreas to PEI and diabetes.

## Acknowledgements

We are grateful for the work of the TRM laboratory at Newcastle University including input from Caitlin Brack and Bethany Hunter. We are especially thankful to all donors and their families.

## Financial declaration

This project was supported by the Cystic Fibrosis Trust SRC 019 and the MRC (Quality and Safety in Organ Donation Tissue Bank – Expansion to include Pancreas/Islets, Heart and Lungs) (MR/R014132/1). This project was supported further by a PhD studentship provided by Newcastle University.

## Author contributions

JAMS, NK and YAS designed the research studies and wrote the original manuscript. YAS, DT, GK, SJR, LR, MP, RC, RM, NK performed experiments and/or acquired data. JAMS, SJR, CF, NK supervised projects. ND and MHS performed tissue sampling for QUOD PANC. All authors have reviewed and edited the manuscript approving of the final version.

## Data availability

Data are available upon reasonable request.

## Ethics Approval and Consent to Participate

Use of human tissue was approved by the responsible ethical review boards.

