## Supplementary material for "Morpho-functional timeline of progressive cystic fibrosis pancreatic exocrine and endocrine pathology derived from semi-quantitative scoring and AI-driven quantitative image analysis"

**Supplementary Table S1: CF donors**

| Case Number | Biobank | Age | Sex | Number of blocks | CF Pattern | Stain |
|---|---|---|---|---|---|---|
| 1 | Klöppel | Premature | Female | 1 | Pattern 1 | H&E, SRFG, CGA / CD31, INS / PP, GLU / SST, α-SMA, CD45 |
| 2 | Klöppel | Premature | Female | 3 | Pattern 1 | H&E, SRFG, CGA / CD31, INS / PP, GLU / SST, α-SMA, CD45, CD68 (2 / 3 blocks) |
| 3 | Klöppel | 2 days | Male | 1 | Pattern 1 | H&E, SRFG, CGA / CD31, INS / PP, GLU / SST, α-SMA, CD45 |
| 4 | EADB | 3 days | NA | 1 | Pattern 1 | H&E, SRFG, CGA / CD31 |
| 5 | EADB | 10 days | NA | 1 | Pattern 1 | H&E, SRFG, CGA / CD31 |
| 6 | EADB | 3 months | NA | 1 | Pattern 1 | H&E, SRFG, CGA / CD31 |
| 7 | EADB | 3 months | NA | 1 | Pattern 1 | H&E, SRFG, CGA / CD31 |
| 8 | EADB | 4 months | NA | 1 | Pattern 1 | H&E, SRFG, CGA / CD31 |
| 9 | Klöppel | 4 months | Male | 2 | Pattern 1 | H&E, SRFG, CGA / CD31, INS / PP, GLU / SST, α-SMA, CD45, CD68 (1 / 2 blocks) |
| 10 | EADB | 6 months | NA | 1 | Pattern 1 | H&E, SRFG, CGA / CD31 |
| 11 | EADB | 2 years | NA | 1 | Pattern 1 | H&E, SRFG, CGA / CD31 |
| 12 | EADB | 2 years | NA | 1 | Pattern 1 | H&E, SRFG, CGA / CD31 |
| 13 | EADB | 3 years | NA | 1 | Pattern 1 | H&E, SRFG, CGA / CD31 |
| 14 | EADB | 3 years | NA | 1 | Pattern 2 | H&E, SRFG, CGA / CD31 |
| 15 | EADB | 4 years | NA | 1 | Pattern 1 | H&E, SRFG, CGA / CD31 |
| 16 | EADB | 4 years | NA | 1 | Pattern 1 | H&E, SRFG, CGA / CD31 |
| 17 | EADB | 4 years | NA | 1 | Pattern 2 | H&E, SRFG, CGA / CD31 |
| 18 | EADB | 7 years | NA | 1 | Pattern 2 | H&E, SRFG, CGA / CD31 |
| 19 | EADB | 7 years | NA | 1 | Pattern 3 | H&E, SRFG, CGA / CD31 |

| | | | | | | |
|---|---|---|---|---|---|---|
| 20 | Klöppel | 7 years | Male | 1 | Pattern 2 | H&E, SRFG, CGA / CD31, INS / PP, GLU / SST, α-SMA, CD45 |
| 21 | EADB | 12 years | NA | 1 | Pattern 3 | H&E, SRFG, CGA / CD31 |
| 22 | Klöppel | 13 years | Female | 7 | Pattern 3 (5 blocks) Pattern 2 (2 blocks) | H&E, SRFG, CGA / CD31, INS / PP, GLU / SST, α-SMA, CD45, CD68 (6 / 7 blocks) |
| 23 | EADB | 14 years | NA | 1 | Pattern 2 | H&E, SRFG, CGA / CD31 |
| 24 | EADB | 14 years | NA | 1 | Pattern 2 | H&E, SRFG, CGA / CD31 |
| 25 | Klöppel | 14 years | Female | 1 | Pattern 2 | H&E, SRFG, CGA / CD31, INS / PP, GLU / SST, α-SMA, CD45 |
| 26 | EADB | 19 years | NA | 1 | Pattern 3 | H&E, SRFG, CGA / CD31 |
| 27 | Klöppel | 19 years | Male | 2 | Pattern 2 | H&E, SRFG, CGA / CD31, INS / PP, GLU / SST, α-SMA, CD45, CD68 (1 / 2 blocks) |
| 28 | Klöppel | 27 years | Male | 3 | Pattern 3 | H&E, SRFG, CGA / CD31, INS / PP, GLU / SST, α-SMA, CD45, CD68 (2 / 3 blocks) |
| 29 | Klöppel | 27 years | Male | 6 | Pattern 2 (4 blocks) Pattern 3 (2 blocks) | H&E, SRFG, CGA / CD31, INS / PP, GLU / SST, α-SMA, CD45, CD68 (5 / 6 blocks) |
| | | Mean age: 6.31 years | | | | |

Pattern 1= Fibrotic, Pattern 2= Fibrotic and lipotic, Pattern 3= Lipoatrophic. Not applicable (NA). EADB cohort: Exeter Archival Diabetes Biobank. Klöppel cohort: Provided by Prof. Günter Klöppel.

**Supplementary Table S2: Control donors**

| Case No. | Biobank | Age | Sex | Stain |
|---|---|---|---|---|
| 1 | EADB | 0 | NA | H&E |
| 2 | EADB | 7 days | NA | H&E |
| 3 | EADB | 7 days | NA | H&E |
| 4 | EADB | 7 days | NA | H&E |
| 5 | EADB | 7 days | NA | H&E |
| 6 | EADB | 13 days | NA | H&E |
| 7 | EADB | 3 weeks | NA | H&E |
| 8 | EADB | 3 weeks | NA | H&E |
| 9 | EADB | 3 weeks | NA | H&E |
| 10 | EADB | 3 weeks | NA | H&E |
| 11 | EADB | 6.2 weeks | NA | H&E |
| 12 | EADB | 4 months | NA | H&E |
| 13 | EADB | 8 months | NA | H&E |
| 14 | EADB | 1 year | NA | H&E |
| 15 | EADB | 2 years | NA | H&E |
| 16 | EADB | 2 years | NA | H&E |
| 17 | EADB | 2 years | NA | H&E |
| 18 | EADB | 2 years | NA | H&E |
| 19 | EADB | 2 years | NA | H&E |
| 20 | EADB | 2 years | NA | H&E |
| 21 | EADB | 2 years | NA | H&E |
| 22 | EADB | 2.5 years | NA | H&E |
| 23 | EADB | 3 years | NA | H&E |
| 24 | EADB | 3 years | NA | H&E |
| 25 | EADB | 3 years | NA | H&E |
| 26 | EADB | 4 years | NA | H&E |
| 27 | EADB | 4 years | NA | H&E |
| 28 | EADB | 5 years | NA | H&E |
| 29 | EADB | 5 years | NA | H&E |
| 30 | EADB | 5 years | NA | H&E |

| | | | | |
|---|---|---|---|---|
| 31 | QUOD | 6 years | Female | H&E, SRFG, CGA / CD31, CD45, INS / PP, GLU / SST, α-SMA |
| 32 | EADB | 6 years | NA | H&E |
| 33 | EADB | 6 years | NA | H&E |
| 34 | EADB | 6 years | NA | H&E |
| 35 | EADB | 6 years | NA | H&E |
| 36 | EADB | 7 years | NA | H&E |
| 37 | EADB | 7 years | NA | H&E |
| 38 | EADB | 7 years | NA | H&E |
| 39 | EADB | 7 years | NA | H&E |
| 40 | EADB | 8 years | NA | H&E |
| 41 | EADB | 8 years | NA | H&E |
| 42 | EADB | 9 years | NA | H&E |
| 43 | EADB | 9 years | NA | H&E |
| 44 | EADB | 10 years | NA | H&E |
| 45 | EADB | 10 years | NA | H&E |
| 46 | EADB | 10 years | NA | H&E |
| 47 | EADB | 10 years | NA | H&E |
| 48 | EADB | 12 years | NA | H&E |
| 49 | EADB | 12 years | NA | H&E |
| 50 | QUOD | 13 years | Male | H&E, SRFG, CGA / CD31, CD45, INS / PP, GLU / SST, α-SMA |
| 51 | QUOD | 18 years | Female | H&E, SRFG, CGA / CD31, CD45, INS / PP, GLU / SST, α-SMA |
| 52 | QUOD | 18 years | Female | H&E, SRFG, CGA / CD31, CD45, INS / PP, GLU / SST, α-SMA |
| 53 | QUOD | 19 years | Male | H&E, SRFG, CGA / CD31, CD45, INS / PP, GLU / SST, α-SMA |
| 54 | QUOD | 24 years | Male | H&E, SRFG, CGA / CD31, CD45, INS / PP, GLU / SST, α-SMA |
| 55 | QUOD | 27 years | Male | H&E, SRFG, CGA / CD31, CD45, INS / PP, GLU / SST, α-SMA |

| | | | | |
|---|---|---|---|---|
| 56 | QUOD | 27 years | Male | H&E, SRFG, CGA / CD31, CD45, INS / PP, GLU / SST, α-SMA |
| 57 | QUOD | 27 years | Male | H&E, SRFG, CGA / CD31, CD45, INS / PP, GLU / SST, α-SMA |
| 58 | QUOD | 29 years | Male | H&E, SRFG, CGA / CD31, CD45, INS / PP, GLU / SST, α-SMA |
| | | Mean age: 7.05 years | | |

EADB cohort: Exeter Archival Diabetes Biobank. QUOD: Quality in Organ Donation.

**Supplementary Table S3: Additional parameters in control deceased organ donors**

| Case No. | Age [years] | Sex | Cause of death | Donor type | Peak CRP [mg/l] | Cardiac and/or respiratory arrest | Major infection |
|---|---|---|---|---|---|---|---|
| 31 | 6 | Female | Hypoxic brain injury - all causes | DBD | 49 | No | no |
| 50 | 13 | Male | Hypoxic brain injury - all causes | DBD | 66 | Yes | unknown |
| 51 | 18 | Female | Hypoxic brain injury - all causes | DBD | 174 | Yes | unknown |
| 52 | 18 | Female | Hypoxic brain injury - all causes | DBD | 204 | Yes | unknown |
| 53 | 19 | Male | Hypoxic brain injury - all causes | DCD | NA | Yes | Pneumonia |
| 54 | 24 | Male | Hypoxic brain injury - all causes | DCD | 3 | Yes | Gram positive coccus |
| 55 | 27 | Male | Intracranial haemorrhage | DBD | 117 | Yes | Bacillus |
| 56 | 27 | Male | Other trauma - accident | DCD | 292 | No | Chest Infection/ pneumonia. |
| 57 | 27 | Female | Intracranial haemorrhage | DBD | 268 | No | no |
| 58 | 29 | Male | Cardiac arrest | DCD | 93 | Yes | unknown |

DBD – Donor after brainstem death, DCD – Donor after circulatory death, NA – Not available, CRP – C-reactive protein

**Supplementary Table S4: Antibody specifications**

| Antibody | Reference | Dilution | Company | RRID |
|---|---|---|---|---|
| Insulin | BSH-2010-100 | 1:1000 | Nordic Biosite, Täby, Sweden | NA |
| Glucagon | EP74 | 1:1500 | Cell Marque, California, USA | NA |
| Pancreatic polypeptide | ab113694 | 1:2000 | Abcam, Cambridge, UK | AB_11156699 |
| Somatostatin | EP130 | 1:300 | Cell Marque, California, USA | NA |
| Chromogranin A | ab254557 | 1:5000 | Abcam, Cambridge, UK | NA |
| CD45 | 2B11+PD7/26 | 1:250 | Dako, Agilent Technologies, Santa Clara, California, USA | NA |
| CD31 | ab182981 | 1:2000 | Abcam, Cambridge, UK | AB_2920881 |
| α-SMA | ab150301 | 1:200 | Abcam, Cambridge, UK | NA |
| CD68 | M0876 | 1:200 | Dako, Agilent Technologies, Santa Clara, USA | NA |

**Supplementary Table S5: Semi-quantitative scoring categories**

| Score | 0 | 1 | 2 | 3 |
|---|---|---|---|---|
| **Ductal lumen dilation** | Non-dilated | Little dilatation sometimes filled with mucus | Dilated and often filled with mucus | Massive dilatation often filled with mucus |
| **Ductal loss** | No ductal loss | Focal ductal loss | Moderate ductal loss | Severe ductal loss |
| **Exocrine pancreas fibrosis** | No abnormal presence of fibrotic tissue | Mild presence of fibrotic tissue | Moderate presence of fibrotic tissue | Massive presence of fibrotic tissue |
| **Acinar atrophy** | No acinar atrophy | <1/3 acini lost | 1/3 – 2/3 acini loss | >2-3 acini loss |
| **Islet remodeling** | No abnormal size or distribution | Some solitary islets surrounded by fibrosis | Many islets are aggregated in complexes | Most islets of variable size and shape are aggregated in complexes |
| **Inflammation** | Scattered inflammatory cells | Mild inflammatory cell infiltration | Moderate inflammatory cell infiltration | Severe inflammation |

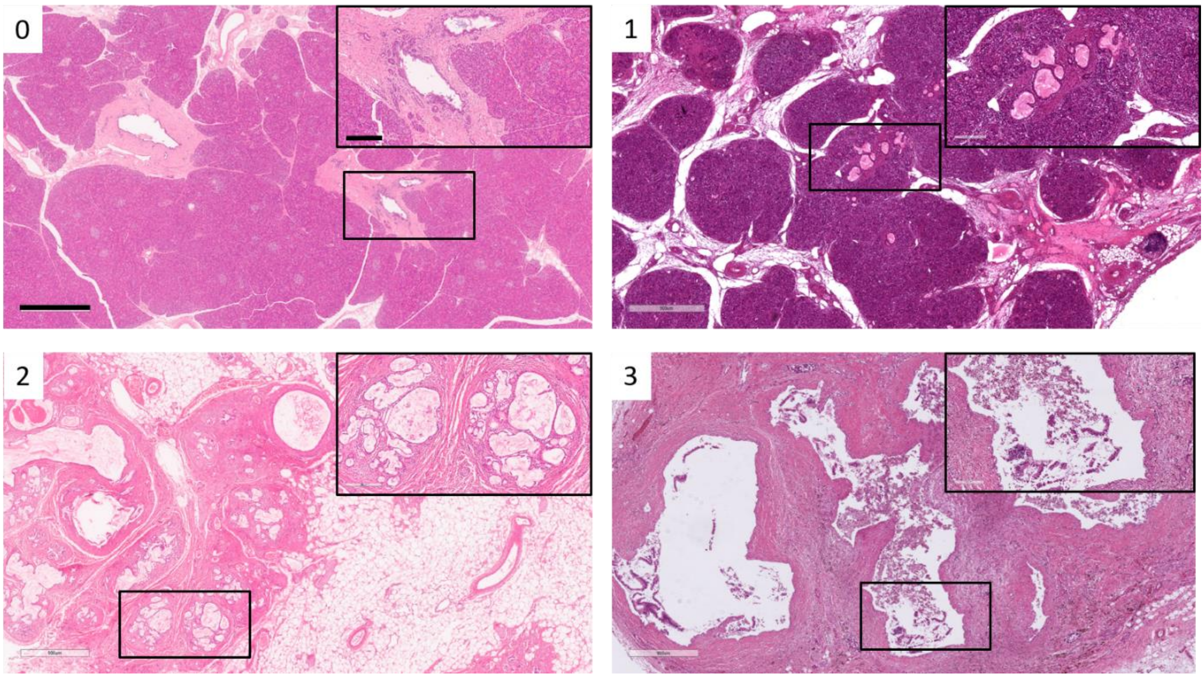


**Supplementary Figure S1: Representative H&E stained control (0) and CF (1-3) pancreatic tissue to exemplify the semi-quantitative scoring system for ductal lumen dilation.** 0: non-dilated; 1: little dilatation sometimes filled with mucus; 2: ducts dilated and often filled with mucus; 3: massive dilatation. Scale bars 900 µm and 200 µm for magnified images (top right corner).

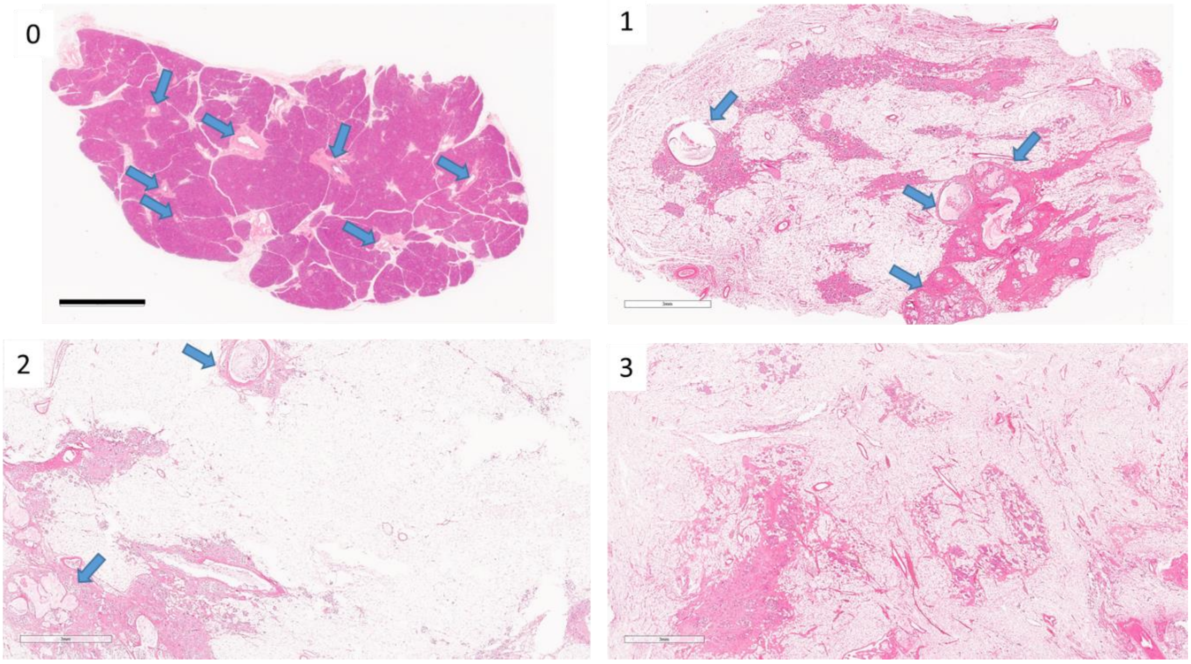


**Supplementary Figure S2: Representative H&E stained control (0) and CF pancreatic tissue (1-3) to exemplify the semi-quantitative scoring system for ductal loss.** 0: no ductal loss; 1: focal loss; 2: moderate loss; 3: severe loss. Blue arrows indicate ducts. Scale bar 3 mm.

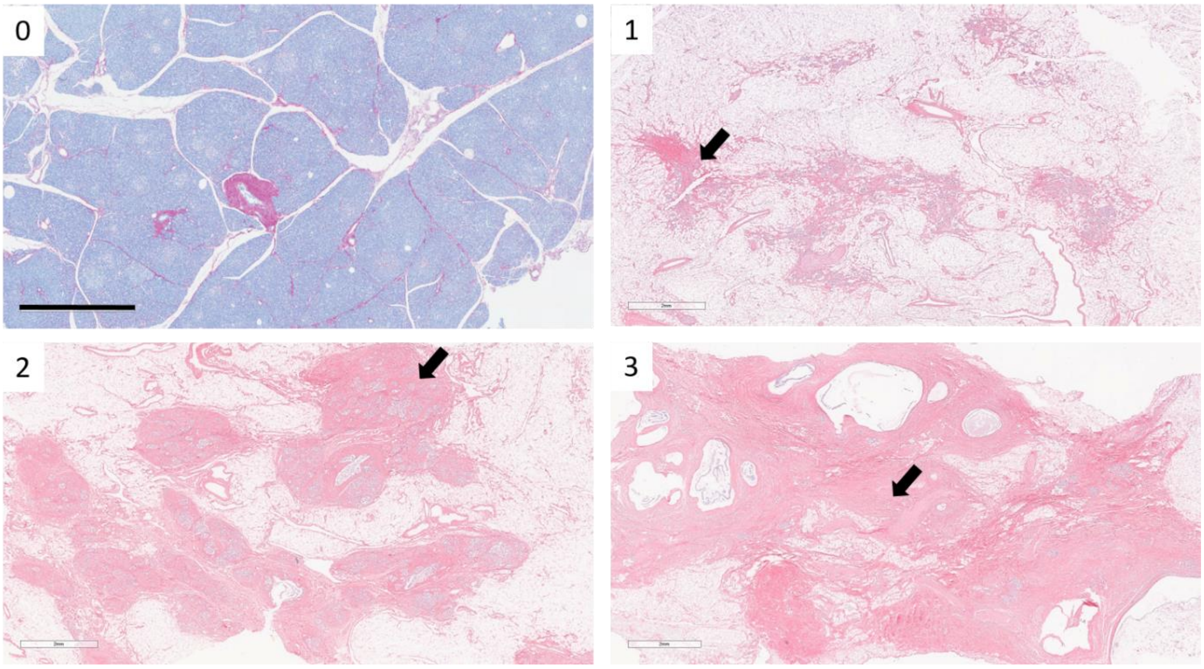


**Supplementary Figure S3: Representative SRFG stained control (0) and CF pancreatic tissue (1-3) to exemplify the semi-quantitative scoring system for exocrine pancreas fibrosis.** 0: no abnormal presence of fibrotic tissue; 1: mild presence of fibrotic tissue; 2: moderate presence of fibrotic tissue; 3: massive presence of fibrotic tissue. Black arrows indicate examples of fibrotic areas. Scale bar 2 mm.

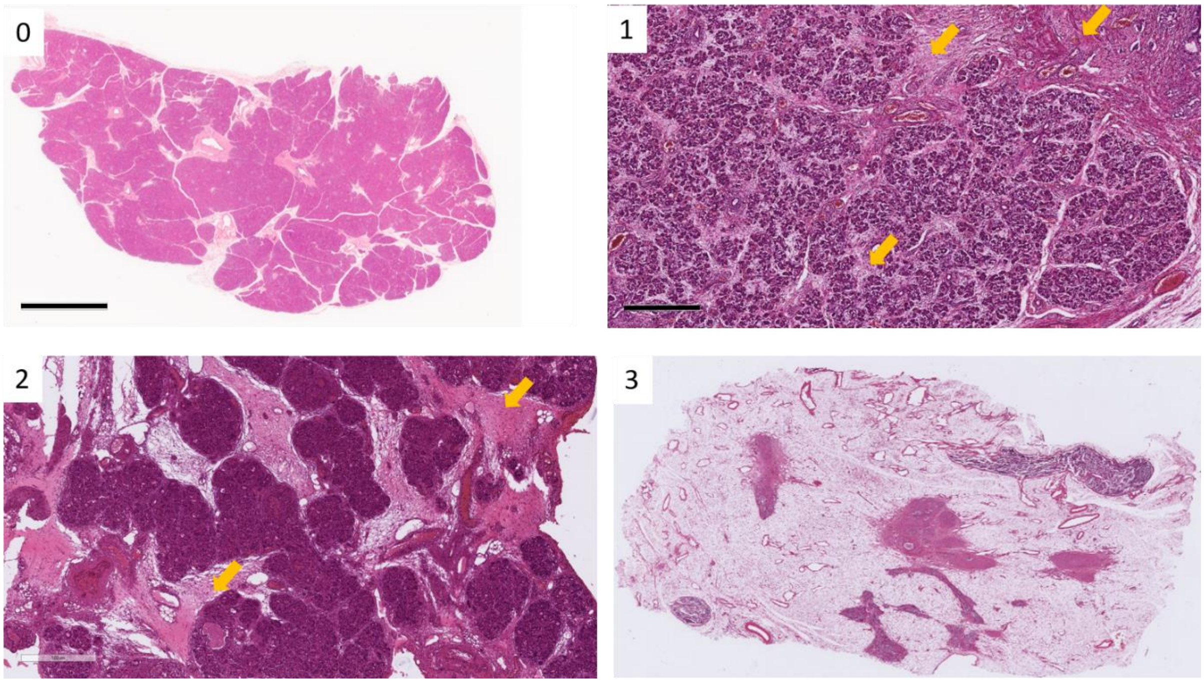


**Supplementary Figure S4: Representative H&E stained control (0) and CF pancreatic tissue (1-3) to exemplify the semi-quantitative scoring system for acinar atrophy.** 0: no acinar atrophy; 1: < 1/3 of acini lost; 2: 1/3 – 2/3 acini loss; 3: >2/3 acini loss. Yellow arrows indicate example areas of acinar atrophy. Scale bar 500 µm (1 &2) and 4 mm (0&3).

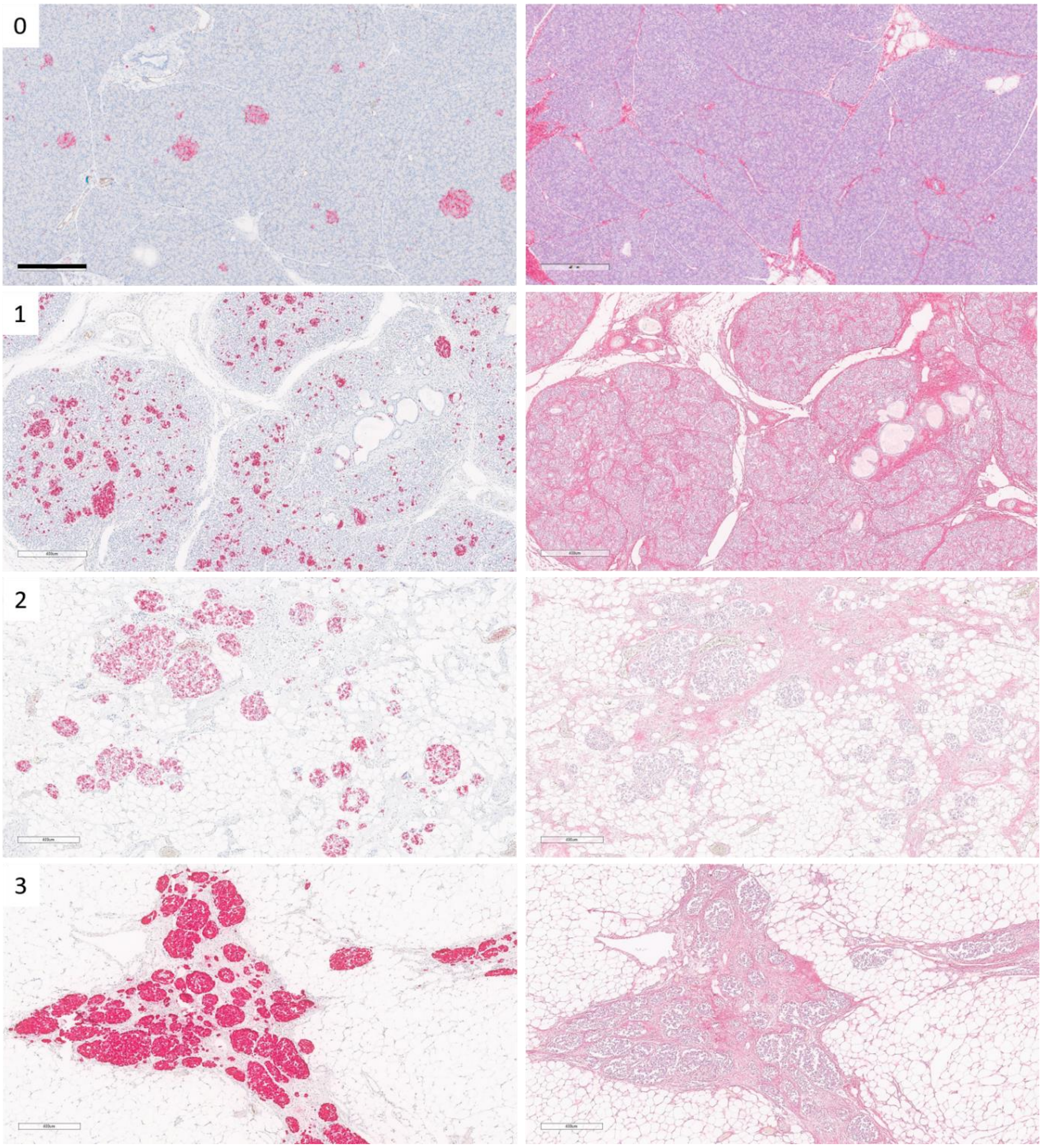


**Supplementary Figure S5: Representative Chromogranin A (left column) and SRFG (right column) stained control (0) and CF pancreatic tissue (1-3) to exemplify the semi-quantitative scoring system for islet remodelling.** 0: no abnormal size or distribution; 1: some solitary islets surrounded by fibrosis; 2: many islets are aggregated in complexes; 3: most islets of variable size and shape are aggregated in complexes. Scale bar 400 µm.

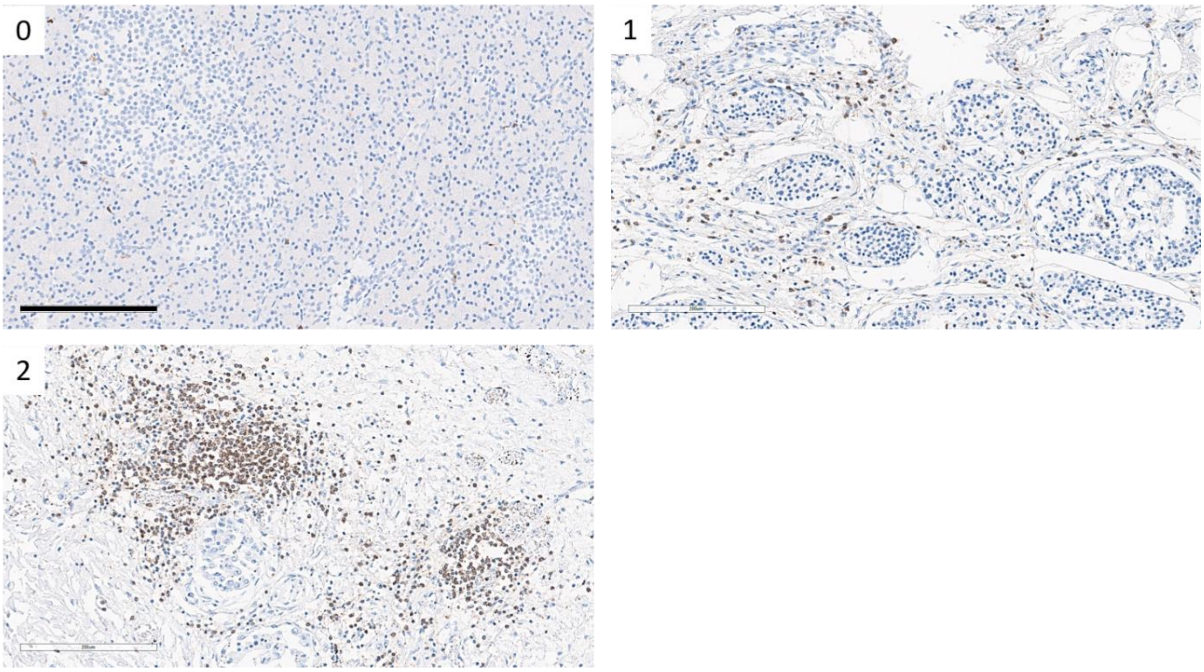


**Supplementary Figure S6**: **Representative CD45 (brown) stained control (0) and CF pancreatic tissue (1-2) to exemplify the semi-quantitative scoring system for inflammation.** 0: scattered inflammatory cells; 1= mild inflammatory cell infiltration; 2= moderate inflammatory cell infiltration; 3= severe inflammation (not present in any samples examined). Scale bar 200 µm.

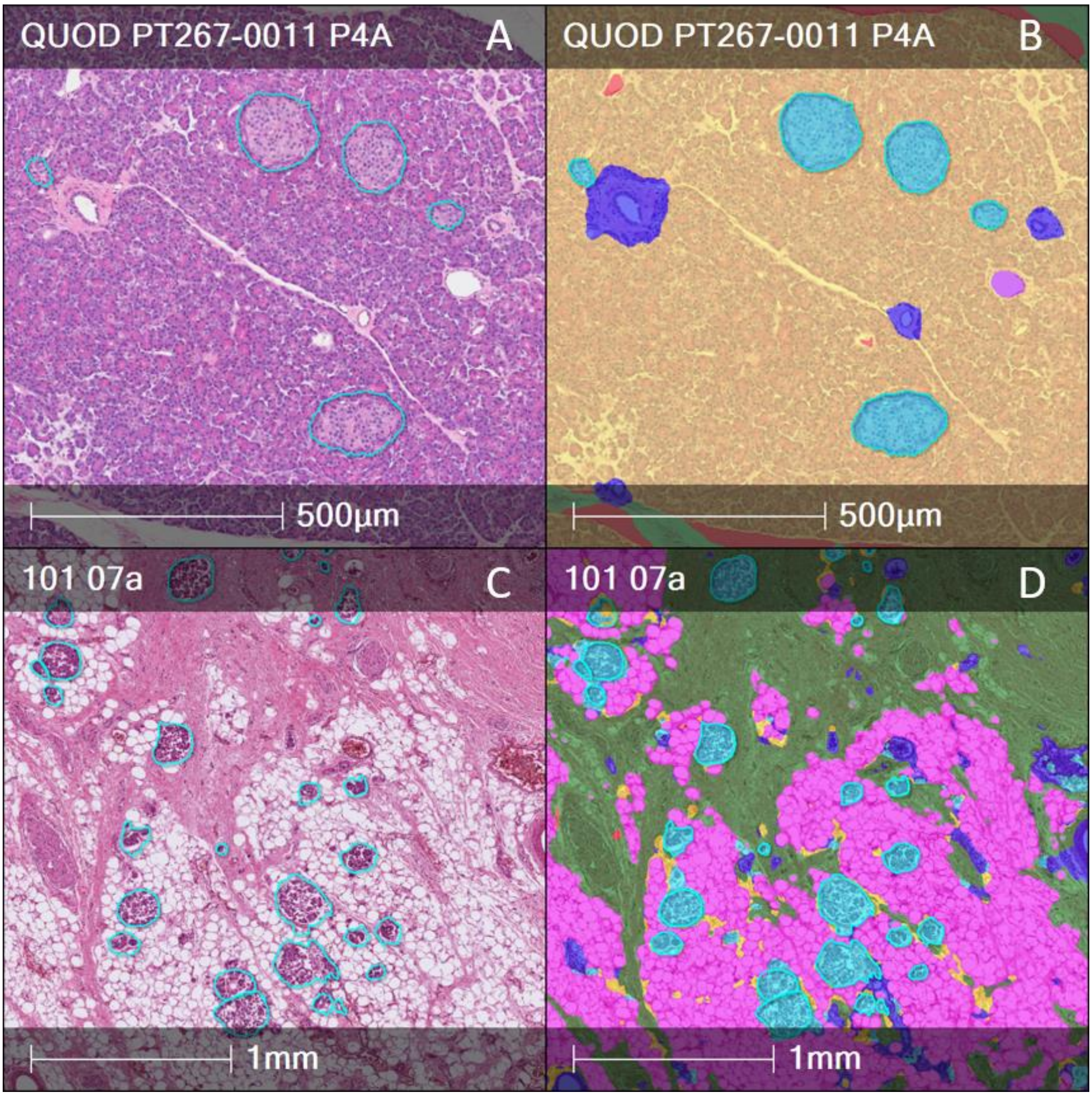


**Supplementary Figure S7: AI classifier illustration of H&E area quantification in a 27 years' old male QUOD control (Case number: 29) (A and B) and a 27 years' old male CF donor (Case number: CF28) (C and D).**

Tissue classifier colours: islets (cyan); ducts (blue); fat (pink); acinar (yellow); background (red); fibrosis (green).

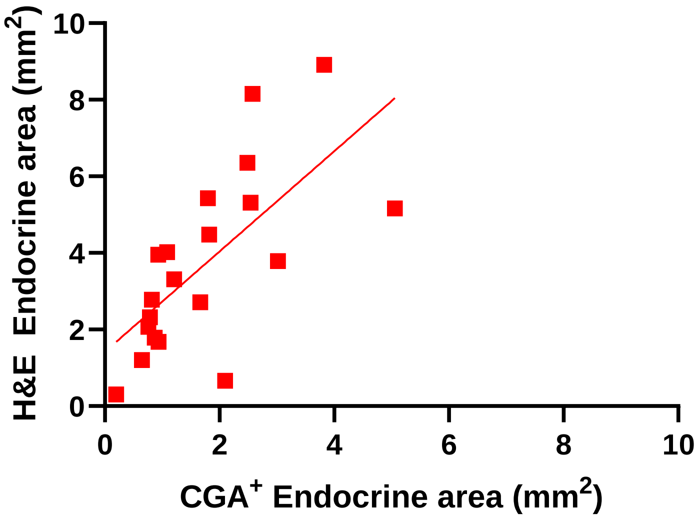


**Supplementary Figure S8: Block-based AI assessment of endocrine area using H&E staining and CGA staining of CF pancreata of different patterns.** Pearson's correlation plot to show correlation between classified endocrine area in CF tissue (20 blocks) using H&E staining and CGA staining to check comparability, $r^2$= 0.465, p=0.0009.

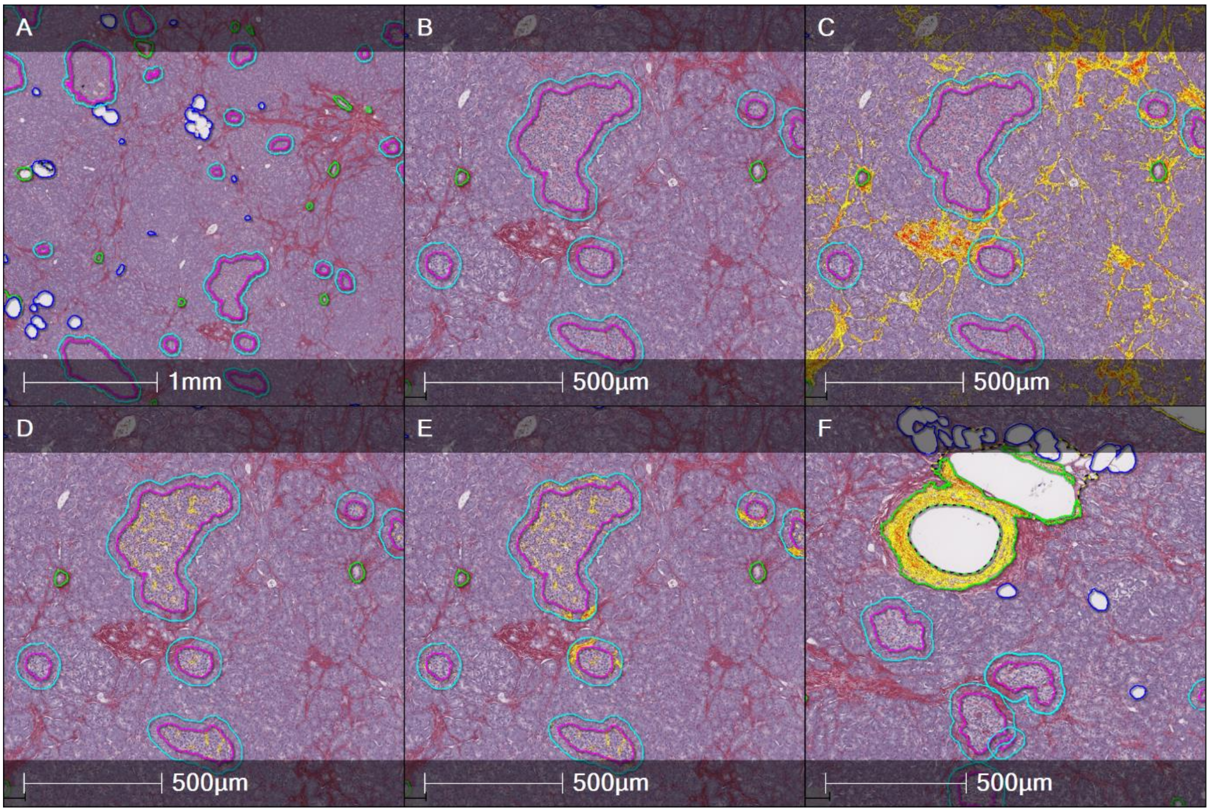


**Supplementary Figure S9: AI classifier illustration of SRFG-positive area (yellow/orange labelling) quantified in different compartments in a 52 years' old male QUOD control (not included in the current study cohort).**

**(A)** Tissue with classifier: fat (blue); islet (pink); peri-islet/ islet (cyan); duct and other (green). **(B)** Tissue classifier – higher magnification. **(C)** SRFG-positive area quantification within acinar compartment. **(D)** SRFG-positive area quantification within islet. **(E)** SRFG-positive area quantification within peri-islet/ islet. Peri-islet SRFG-positive area was calculated by subtracting islet area (pink) from peri-islet/islet area (cyan). **(F)** SRFG-positive area quantification within duct and other (blood vessels).

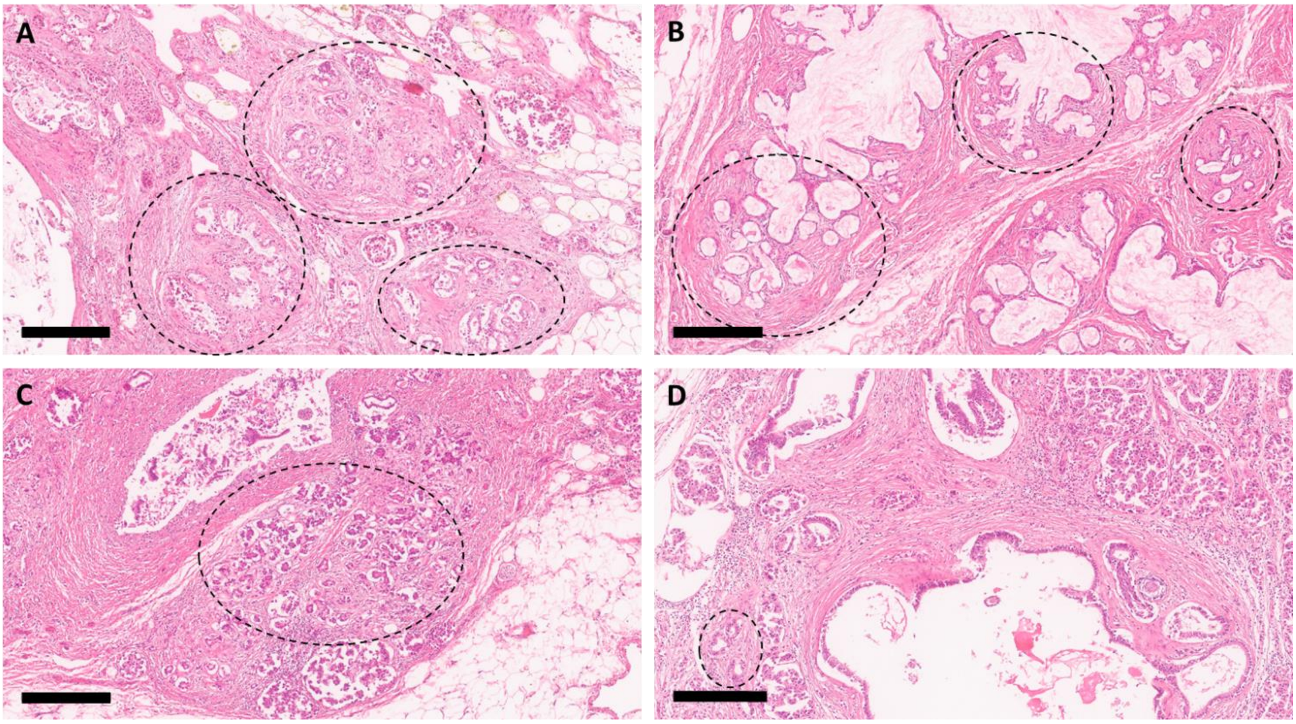


**Supplementary Figure S10:** H&E staining of CF Case number 28 (A), 22 (B), 27 (C), and 29 (D). Dashed circles show small residual ducts surrounded by fibrosis. Scale bars 300 µm.

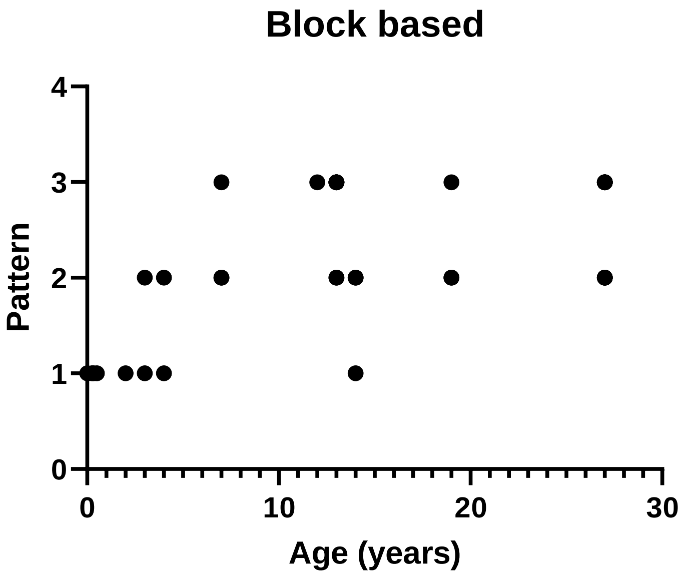


**Supplementary Figure S11: Block based pattern analysis of donors with CF vs age.** Spearman correlation coefficient r=0.7156, p<0.0001.

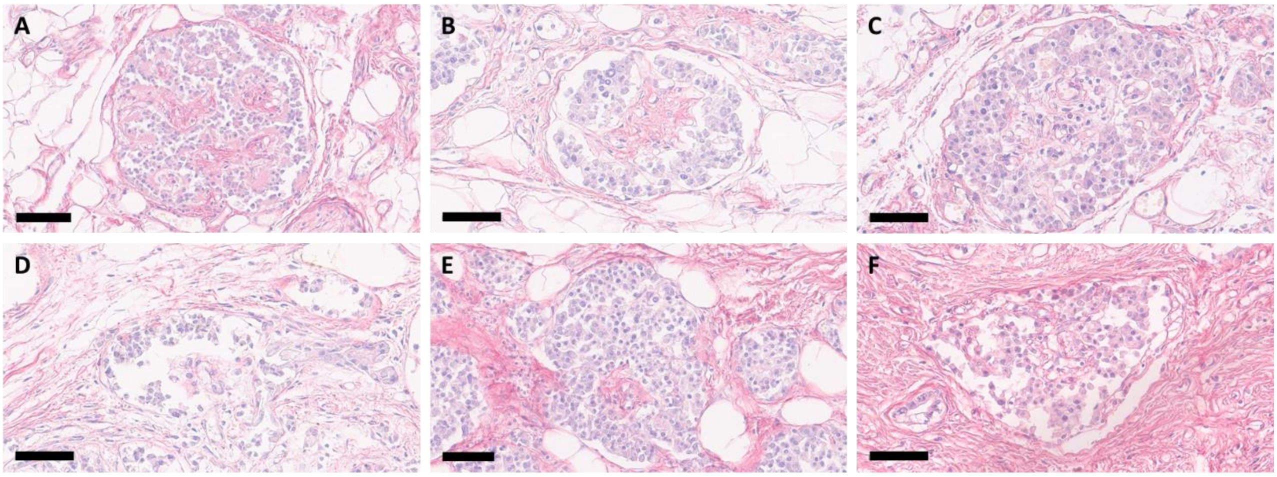


**Supplementary Figure 12:** SRFG staining of CF Case number 22 (A-C), 28 (D), 29 (E) and 27 (F) showing intra-islet and peri-islet fibrosis. Scale bars: 60 µm (B, C, D, F), 70 µm (E), 80 µm (A).

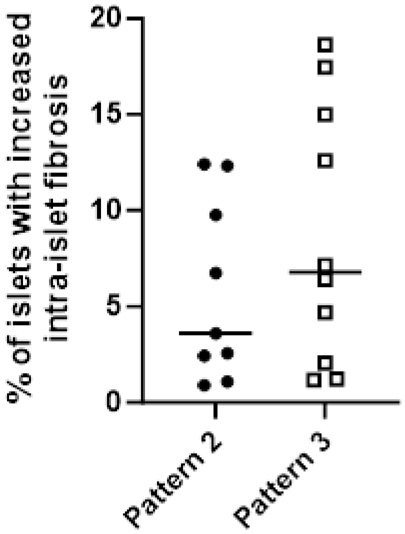


**Supplementary Figure 13: Manual quantification of the percentage of islets with significant fibrosis extending beyond the intra-islet perivascular region in CF Pattern 2 and CF Pattern 3.** No significant difference between the patterns was observed.

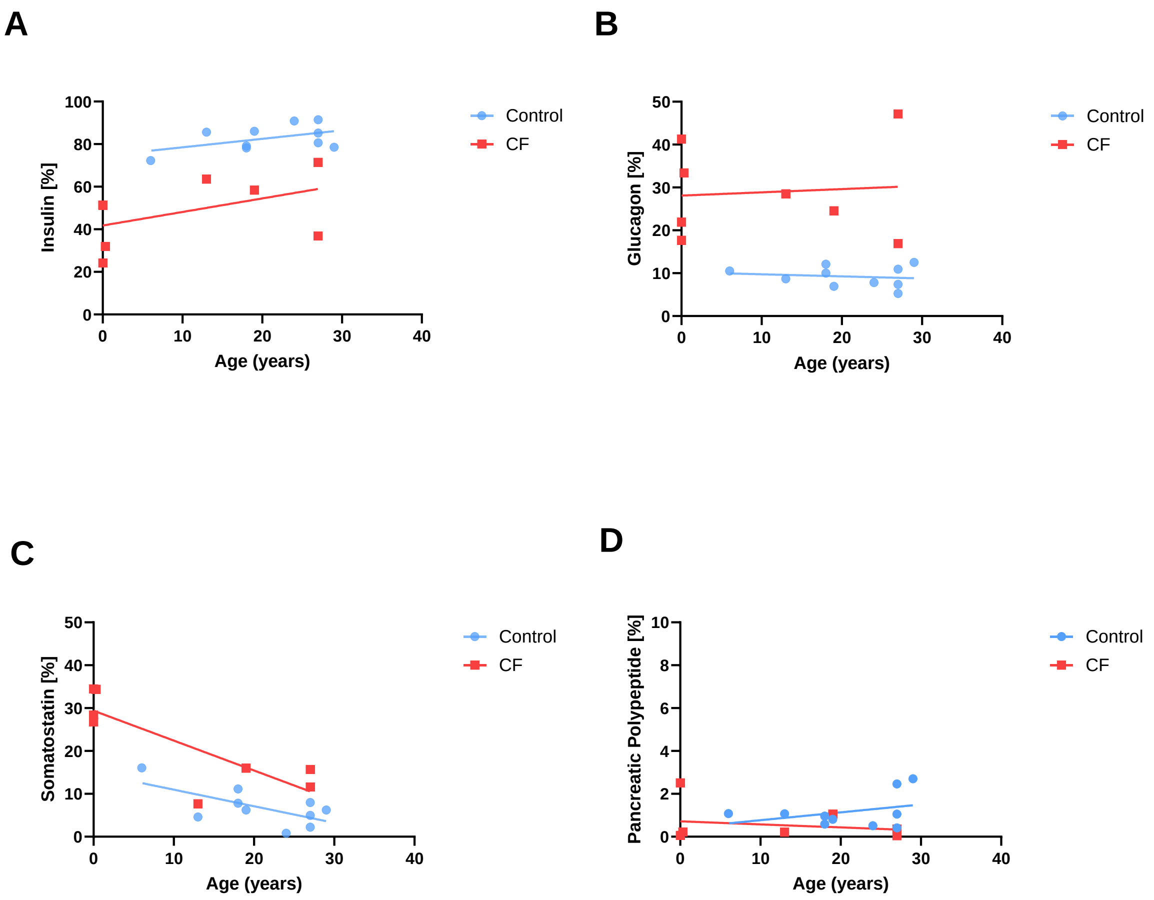


**Supplementary Figure S14: Islet hormone distribution in CF and non-CF pancreata**. Percentage insulin (A), glucagon (B), somatostatin (C) and pancreatic polypeptide (D) PA vs donor age in control (blue) and CF (red) donors. For donors with multiple blocks mean values were used.

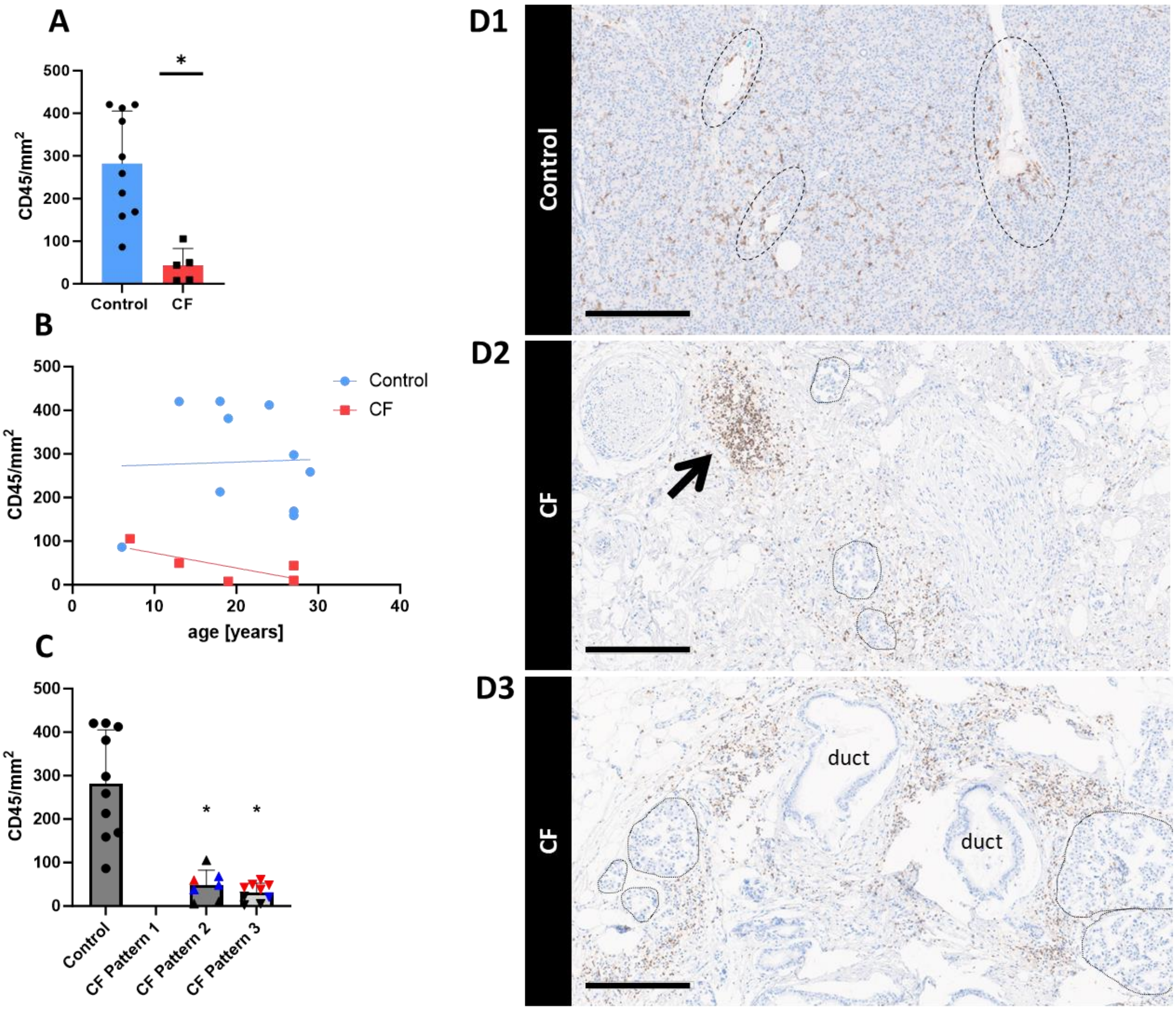


**Supplementary Figure S15: Leucocyte distribution in control and CF pancreata.** (A) Donor based AI analysis of CD45-positive leucocyte count/mm$^2$ in control (blue) and CF (red) donors. (B) Donor-based AI analysis of CD45-positive lymphocyte count/mm$^2$ vs age of control (blue circles) and CF (red squares) donors. (C) Pattern-based AI analysis of CD45-positive leucocyte count/mm$^2$ in control and CF pancreata. Donor-based statistical comparisons were carried out using unpaired Student's T-test and block-based analyses were conducted using linear mixed effect model (LMEM) with pancreata pattern set as fixed effect, donor as random effect and Bonferroni post hoc test. Graphs show the mean±SD. $p<0.05$ was considered significant. (*) indicates significant difference compared to control. Red triangles represent Case 22 and blue triangles represent Case 29: both donors have multiple blocks of different patterns. (A, B) control: n=10 donors, CF n=5 donors. (C) control n=10 blocks, CF Pattern 1: n=0 blocks, CF Pattern 2 n=7 blocks, CF Pattern 3 n=9 blocks. (D1-D3) Example pictures of CD45 immunostaining (brown) of control (D1, donor Number 50) and CF pancreata (D2, D3; donor Number 29). Dashed lines highlight vessels, solid lines highlight islets, arrow indicates inflammatory focus. Scale bars 300 µm.

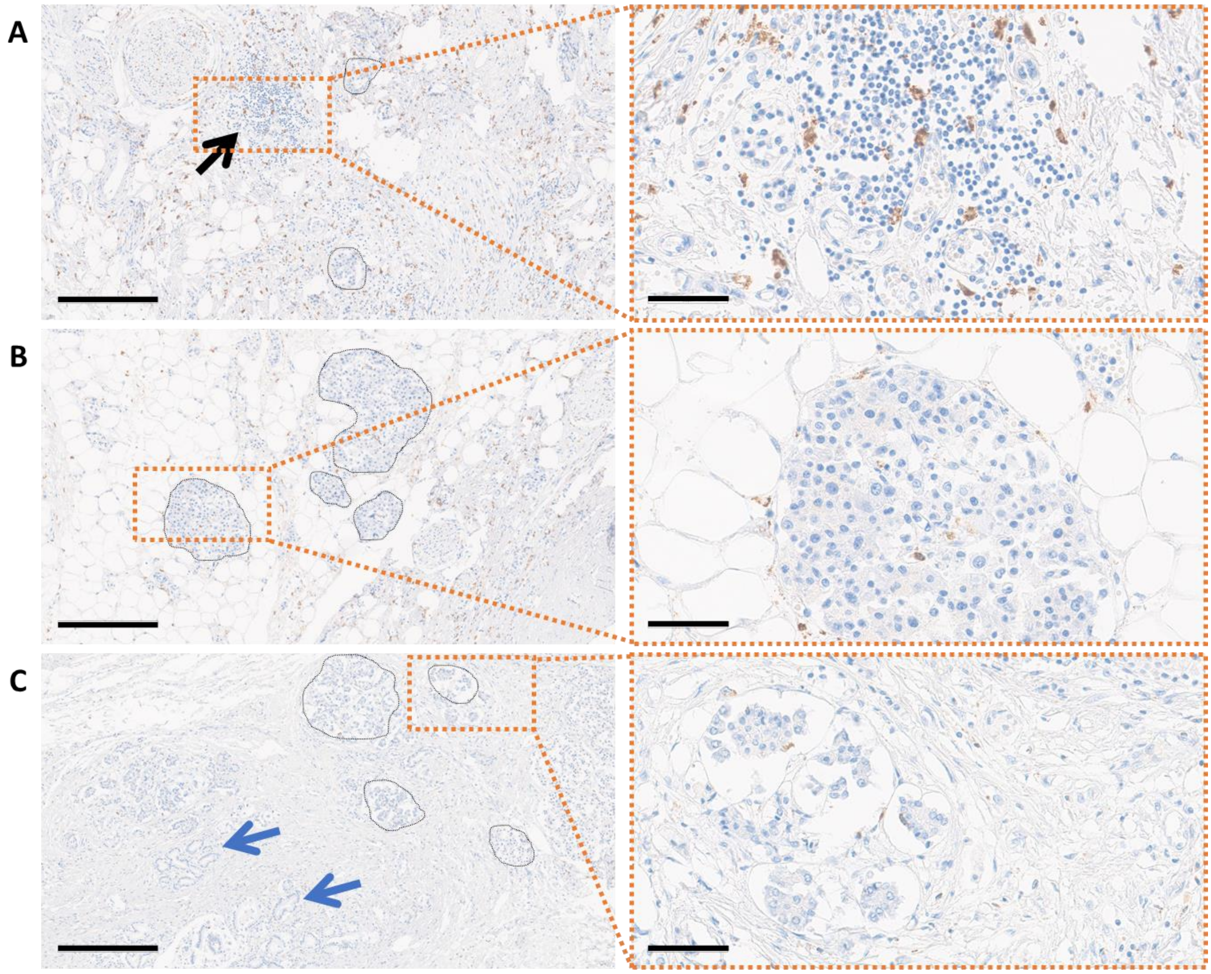


**Supplementary Figure S16. CF pancreas stained for CD68 (brown).** A, B: different blocks from donor Number 29. C: donor Number 27. Scale bars: 300 µm left column, 60 µm right column. A shows a corresponding area to the CD45 staining in Supplementary Figure S15 D2 highlighting macrophages in the lymphocytic focus. B and C show peri- and intra-islet macrophages. Blue arrows in C indicate areas of small ducts.
